# Functional impact of rare variants and sex across the X-chromosome and autosomes

**DOI:** 10.1101/2025.01.23.634570

**Authors:** Rachel A. Ungar, Taibo Li, Nikolai G. Vetr, Nicole Ersaro, Alexis Battle, Stephen B. Montgomery

## Abstract

The human X-chromosome contains hundreds of genes and has well-established impacts on sex differences and traits. However, the X-chromosome is often excluded from many genetic analyses, limiting broader understanding of variant effects. In particular, the functional impact of rare variants on the X-chromosome is understudied. To investigate functional rare variants on the X-chromosome, we use observations of outlier gene expression from GTEx consortium data. We show outlier genes are enriched for having nearby rare variants on the X-chromosome, and this enrichment is stronger for males. Using the RIVER model, we identified 753 rare variants in 449 genes predicted to have functional differences between males and females. We examined the pharmacogenetic implications of these variants and observed that 25% of drugs with a known sex difference in adverse drug reactions were connected to genes that contained a sex-biased rare variant. We further identify that sex-biased rare variants preferentially impact transcription factors with predicted sex-differential binding, such as the XIST-modulated SIX1. Combined, our study investigates functional rare variants on the X-chromosome, and further details how sex-stratification of variant effect prediction improves identification of rare variants with predicted sex-biased effects, transcription factor biology, and pharmacogenomic impacts.

## Introduction

The exclusion of the X-chromosome in genetic studies remains prevalent despite its role in sex determination and human traits. Only ¼ of current GWAS include the X-chromosome, yet it contains 800 genes, over 60% of which have been linked to genetic disorders (Leitão et al. 2022; Sun et al. 2023). Unlike other chromosomes, the X-chromosome differs in copy number between males and females and is also subject to aneuploidy, with up to five copies being reported in humans (Rogol 2020; Khramtsova et al. 2019). The difference in copy number between males and females has been suggested as a potential mechanism for differences in traits and disease between the sexes (Khramtsova et al. 2019).

Sex-biased genes are located across both the autosomes and X-chromosome, but a larger role has been implicated for these genes on the X-chromosome (Oliva et al. 2020; Flynn et al. 2021). Across metazoans, 30-60% of genes are estimated to be sex biased and approximately 37% of human genes are sex-biased in at least one tissue, although with predominantly small effect sizes (Albritton et al. 2014; Oliva et al. 2020). Biological sex can lead to differences in disease risk and even impact drug efficacy and susceptibility to adverse drug reactions (Brabete et al. 2022; Fisher et al. 2022; Khramtsova et al. 2019).

Even among genetic studies that consider sex and the X-chromosome, most have been restricted to common variants. For example, when the X-chromosome is included in analyses of molecular traits like gene expression, a small number of expression quantitative trait loci (eQTLs) are often discovered due to unique selective pressures on the X-chromosome (Le Guen et al. 2021; Gorlov and Amos 2023). Historically few sex-biased eQTLs have been discovered from bulk RNA-seq, likely due to power issues (Oliva et al. 2020; Kukurba et al. 2016; Porcu et al. 2022). As eQTL studies focus on common variants, there remains an ongoing gap in understanding how low-frequency or rare variants may contribute to sex differences in gene expression and, ultimately, complex traits and diseases.

Rare variants, by definition, each occur in few people in the general population (allele frequency < 1%). However, collectively they are common with singletons, alleles seen in just one individual, the most abundant and impactful class of variants (Karczewski et al. 2020; Glassberg et al. 2019). Despite their importance, the study of rare variants cannot easily follow traditional common-variant approaches, such as genome-wide association studies (GWAS), as both multiple-testing burden and sample size required for adequate power impede their analysis. In lieu of association testing, understanding the functional properties of impactful rare variants allows for effective prediction of a functional effect. Prior studies have employed an outlier-based approach to identify functional properties of rare variants impacting extreme gene expression (Bonder et al. 2021; Brechtmann et al. 2018; Frésard et al. 2019; Li et al. 2023; Smail et al. 2022; Ferraro et al. 2020; Li et al. 2017). This approach is analogous to efforts to study extreme phenotypes in population samples to enrich for impactful rare variants (Li et al. 2011; Barnett et al. 2013). However, it has yet to be applied to the X-chromosome and across sexes where dosage and sex-specific regulation can impact discovery.

In our study, we evaluated outlier-associated rare variants on the X-chromosome, and further applied a sex-stratified approach genome-wide across the X-chromosome and the autosomes. Sex-stratification has repeatedly been shown to impact discovery in genomic analyses (Merikangas and Almasy 2020). Despite advances in machine learning and AI-based variant effect prediction, sex-specific datasets have received limited attention. The same applies to rare variant prioritization methods like RIVER that integrate molecular outlier data (Li et al. 2017). Using the Genotype Tissue Expression (GTEx) consortium data, we detected multi-tissue outliers on the X-chromosome, investigated the properties of outlier-enriched rare variants, and the impact of sex-stratification on training and prediction. We subsequently sex-stratified GTEx data, trained and applied the RIVER model to prioritize rare variants with sex-specific functional effects, and examined their potential interactions with drugs and sex-specific transcription factors. Combined, this work highlights rare variant effects on the X-chromosome and the value of sex-informed analyses of variant effects.

## Methods

### Variant filtering and annotation

Variants were obtained from the GTEx v8 dataset (THE GTEX CONSORTIUM 2020). Allele frequencies were then annotated for each site using gnomAD v3.0 (Quinlan and Hall 2010; Karczewski et al. 2020). Following Ferraro et al., individuals were subset to only include those with high European ancestry (genetic PC1<0 and PC2>0.05) due to the low proportion of other ancestries and subsequent smaller group sizes when conducting sex-stratification by ancestry (Ferraro et al. 2020). We further excluded any individuals with atypical sex chromosomes or sexual development. We considered only rare variants within genic regions and at most +/−5kb from a gene. We restricted our analysis to lncRNA and protein-coding genes using GENCODEv26 (Frankish et al. 2019). All variants were subsequently annotated with CADD scores (Rentzsch et al. 2021). For the X-chromosome, we performed extra variant filtering given we retained pseudoautosomal regions. We removed variants in ENCODE blacklisted regions (Amemiya et al. 2019) or with an average genotype quality (GQ) score differing by more than 5 between males and females (Supplementary Figure 14).

### Sex-stratification of GTEx samples

The number of copies of the X-chromosome is one of the components that define the biological sex of an individual. The biological term sex is a multidimensional concept referring to a combination of sex chromosomes, internal and external genitalia, and sex hormones (Ainsworth 2015). In this paper, we used a narrow definition of sex, in which we only considered sex chromosomes, and referred to individuals with XX chromosomes as female and XY chromosomes as male.

For outlier analyses, we stratified our samples into four separate groups: female (F), male (M), combined (all), combined (equal sample size). Sex stratification was done prior to read count filtering. We defined samples as male based on the presence of XY sex chromosomes and the GTEx-reported sex, and females based on the presence of XX sex chromosomes and the GTEx-reported sex. As GTEx is male-biased, we first aimed to control for power differences by subsetting the male group to be of equal sample size with the female group (N_F_=N_M_). Subsequently, the combined (all) group was created by taking all samples (N_F_+N_M_) while the combined (equal sample size) group was created to be of equal sample size (½N_F_+½N_M_=N_B_). In the combined (equal sample size group), the individuals with the maximum number of tissues were included to improve subsequent multi-tissue outlier discovery. Null sets were created by shuffling sex labels.

### Expression processing, correction, and outlier calling

TPM values and read counts were obtained from the GTEx v8 dataset (THE GTEX CONSORTIUM 2020). We filtered to 22189 protein-coding and 783 lncRNA genes on the autosomes and X-chromosome.

Expression data was processed following the same methods as Ferraro et al. (Ferraro et al. 2020). Briefly, read counts are filtered such that in the combined group at least 20% of individuals in at least one sex have a TPM>0.1 and read count >6, and 5% of individuals do not have zero read counts. Then, for each group, these counts are log transformed, centered, and scaled to create z-scores. For each tissue and group, PEER factors were calculated (Stegle et al. 2012). For each tissue and group, corrected counts were calculated using a linear model incorporating the top three genetic PCs, top eQTLs, and PEER factors. The residuals of this regression were then transformed into z-scores to be used for the outlier analysis.

Thresholds for multi-tissue outlier discovery were chosen based on parameterizing across a range of z-scores and tissues (Supplementary Figure 15A,B). Multi-tissue outliers were required to be outliers in at least three tissues, and the absolute median z-score greater than 2.5 across all tissues. Any gene-individual pairs identified as a multi-tissue outlier for more than two individuals were removed.

### Rare Variant Relative Risks for Expression Outliers

Enrichment of rare variants in gene expression outliers were calculated in R using epitools (Aragon et al. 2020). For this analysis, rare variants were first collapsed to the gene level to assess the presence of any rare variant which also satisfied any additional annotation criteria. Considered annotations were whether there was a variant with greater than or equal to a CADD threshold (in practice this is 0 or 15) and within a given distance (5 kb) from the gene start and end (Supplemental Figure 15C). If multiple variants passed the annotation and frequency filters, we selected the variant with the lowest minor allele frequency. We subsequently binarized “rare” for an individual-gene pair (whether outlier or non-outlier) depending on different annotation and allele frequency thresholds in our relative risk enrichment analyses.

For relative risk calculations, we assessed the proportion of outliers with a threshold-passing rare variant compared to non-outliers.

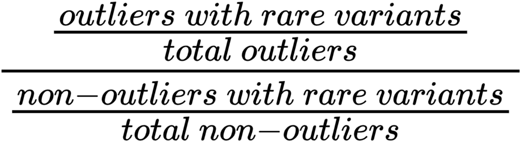

A relative risk score greater than one indicated that there was an enrichment for outliers to have a threshold-passing rare variant compared to non-outliers, a score less than one indicated outliers were depleted of rare variation.

### Sex-Specific Rare Variant Discovery

To assess the functional impact of rare variants on gene expression, we applied RIVER, a Bayesian hierarchical model which integrates genomic annotations and observed outlier effects (Li et al. 2017). For model training, we used a set of N=30 genomic annotations calculated from VEP, CADD, and genomic locations summarized across all rare variants within 10kb of the gene (G layer), and used median z-score across tissues estimated from a balanced set of female and male individuals as observed outlier signals (E layer). Binary outlier threshold was set at p < 0.05 (corresponding to approximate |Z| > 2, --pvalue_threshold=0.05). Approximately 1% of gene-individual pairs reached this threshold, which we used for model evaluation (--pvalue_fraction=0.01). We next separately modeled over- and under-outliers as previous work has identified different categories of rare variants contributing to these signals (Ferraro et al. 2020). To enrich for outlier signals with genetic effects, we filtered out genes which do not have any outlier individuals (defined as |Z| > 2).

For model evaluation, we leveraged pairs of individuals with the same set of rare variants nearby the same gene (“N2 pair”), where we inferred the regulatory status of the second individual based on genomic annotations and observed outlier status of the first individual. Area under the precision-recall (P-R) curve was used as the evaluation metric. Importantly, these N2 pairs were not included in model training. In total, we had 532,975 instances of (gene, individual), of which we had 38,263 N2 pair individuals for evaluation of the model. RIVER scores range between 0 and 1 in magnitude, where 1 indicates a high probability that the variant drives an outlier expression effect. The sign of the score represents the direction of this effect: negative scores mean a variant reduces expression and positive scores mean a variant increases expression.

To estimate sex-specific effects of rare variants, we trained on both sexes, and then conducted predictions within each sex, thereby obtaining posterior probabilities [P(Z | G, E)] of driving outlier expression levels of nearby genes for each variant in each sex. We also trained a model using expression data on the X-chromosome from both sexes, and estimated sex-specific posterior effects similarly. For variants appearing in at least one female and one male, we calculated the median posterior within either females or males to derive the sex-specific posteriors. We defined a variant as sex-biased if the posterior probability in one sex was higher than the posterior probability in the other (at a threshold of 0.2). We further classified sex-specific variants based on direction of effect, where a female under-expression rare variant had a significantly higher posterior for under-expression in females than in males (|female posterior| - |male posterior| > 2 and female posterior < 0), and similarly for the other three categories.

### Annotation of Sex-Specific Rare Variants and Gene Set Enrichment Analysis

For each group of sex-biased rare variants, we assessed their overlap with existing annotations by computing the fraction of variants annotated in each VEP category (McLaren et al. 2016). For each variant type, we evaluated if the number of sex-biased functional rare-variants that had a given annotation as compared to the number without this annotation, was significantly different than that of non-functional rare variants using a Fisher’s Exact Test. This was done for male-biased, female-biased, and non sex-biased functional rare variants, and additionally split for those predicted to increase and reduce expression.

To characterize genes associated with sex-specific rare variants, we took all genes predicted by RIVER to be disrupted by these rare variants. Importantly, for rare variants nearby multiple genes, we only took genes for which RIVER posterior showed sex-specific effects as defined above. We grouped female-biased variants as those with higher posterior in females (female over-expression and male under-expression), and similarly defined male-biased variants as those with higher posteriors in males (male over-expression and female under-expression) at a difference greater than 0.2. We then used the clusterProfiler package in R to assess significantly enriched GO terms in female and male-biased gene sets and performed FDR correction on the resulting p-values (Wu et al. 2021). To assess significance of overlap between genes with a sex-biased rare variant and genes with sex-biased eQTLs reported in GTEx we applied a hypergeometric test (Oliva et al. 2020).

### Pharmacogenetic Annotation and Analysis

Genes with sex-biased rare variants predicted by RIVER were linked to drugs using the Drug Gene Interaction database (Cannon et al. 2024). A list of drugs with adverse drug reactions was obtained from the Table of Pharmacogenomic Biomarkers in Drug Labeling on the FDA Center for Drug Evaluation and Research website (Center for Drug Evaluation and Research 2024). A list of sex-biased adverse drug reactions was obtained from Zucker et al. (Zucker and Prendergast 2020). The relative risk of the proportion of sex-biased rare variants within a given database as compared to non-biased functional rare variants was evaluated using the epitab function using the Wald test from the epitools package (Aragon et al. 2020).

### Sex-Specific Transcription Factor Network Analysis

We performed two analyses to investigate the relationship between prioritized rare variants and transcription factor binding. First, we performed a genome-wide motif enrichment analysis using HOMER around 100 bp up and downstream of each class of variants, setting `-size given` and masking repeat regions with all other parameters as default (Heinz et al. 2010). We obtained the top enriched motifs where the fraction of motifs overlapping the genomic regions centered on sex-specific rare variants are significantly higher than that overlapping with the background, and highlighted top enriched motifs ranked by enrichment p-value. Next, for all rare variants within annotated regulatory regions, we estimated disruption of transcription factor binding using FABIAN-variant, which leverages a large collection of motif databases to model alterations in TF binding between reference and alternative alleles, across all transcription factors in the database (Steinhaus et al. 2022). For each variant and each TF, we summarized predicted effects across all models to obtain final effect estimates. To investigate sex-specific TF effect on target genes, we designed metric to combine predicted rare variant effect on TF binding and the posterior estimates of a rare variant underlying expression outlier signals of a nearby gene, where for TF i and gene j, we calculate the regulatory score across SNPs (v) in the 10kb window as follows

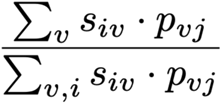

Where s_iv is the predicted disruption of TF i’s motif by variant v, and p_vj is the RIVER posterior of variant v on gene j. The denominator is in place to compare relative regulatory effects of all TFs for each gene.

To control for false positive discovery of sex-specific variants, we randomly permuted sex labels in z-score calculation and outlier calling five times, and recalculated rare variant posteriors by RIVER. We repeated the same motif enrichment and binding disruption analysis, and only kept top TFs with higher enrichment and estimated binding effects than results from all permuted models.

## Results

### Individuals carry multi-tissue gene expression outliers on the X-chromosome

Gene expression outliers can inform likely disease-causing genes and prioritize functional rare variants (Frésard et al. 2019; Ferraro et al. 2020). We focused on gene-individual pairs that were outliers (absolute Z-score > 2.5) in multiple tissues (multi-tissue outliers, see Methods).

We identified multi-tissue outliers on the autosomes and X-chromosome, and selected chromosome 7 (chr7) as a control for outlier discovery given it has a similar number of genes as the X-chromosome. In total, we identified 158 multi-tissue outliers (gene-individual pairs) on the X-chromosome, 245 on chr7, and 4,985 on the autosomes (Figure 1A). This corresponded to an average number of gene expression outliers per individual of 0.21 multi-tissue outliers on the X, 0.33 multi-tissue outliers on chr7, and 6.75 multi-tissue outliers across all the autosomes (Figure 1B).

**Figure 1:**
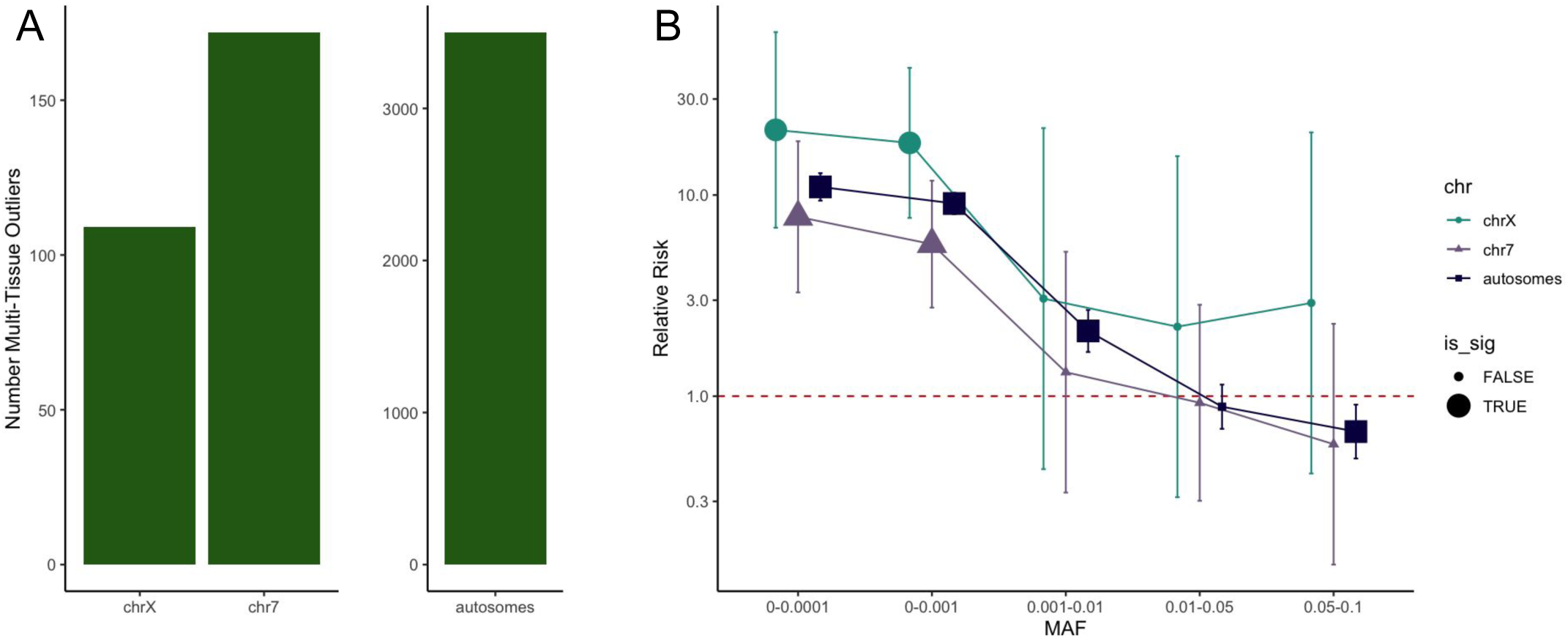
Outliers and rare variants across the autosomes and X-chromosome. a, The number of multi-tissue outliers across the X-chromosome, chromosome 7, and the autosomes. b, The relative risk of outlier having a rare variant within +/− 5kb as compared to a non-outlier across different minor allele frequency (MAF) bins for the X-chromosome, chromosome 7, and autosomes.

Multi-tissue outliers were discovered on the X for 133 unique genes across 135 individuals (18% of total genes tested on the X-chromosome) (Figure 1A). Of these genes, 22 have prior evidence of trait relationships in Open Targets with evidence scores > 0.6. We evaluated if genes harboring outliers reflected other metrics of selective constraint. Overall, there were more over-expression outliers than under-expression outliers on the X-chromosome, consistent with previous studies of autosomes and reflecting higher tolerance for gain-of-expression effects (Supplemental Figure 1A) (Li et al. 2017). The proportion of over-expression outliers on the X-chromosome was significantly higher than in the autosomes (p=0.02).

In gnomAD v3, constraint metrics for genes have been calculated to identify their tolerance for having missense, synonymous, and loss-of-function variants (Chen et al. 2022; Samocha et al. 2014; Lek et al. 2016). Among these metrics, outlier genes were less tolerant of missense variants and over-expression outlier genes showed evidence of being less tolerant of synonymous mutations (Supplemental Figure 1B). However, under-expression outliers were depleted for genes less tolerant of loss-of-function mutations, which could be expected due to the largely non-disease status of the GTEx samples and cohort (Supplemental Figure 1B). We also applied a dosage constraint metric using ANEVA to classify the amount of variation a gene expression is expected to have, and analogously the MoD score to calculate the amount of gene dosage tolerance, where a higher number in each score demonstrates more tolerance (Dong et al. 2023; Mohammadi et al. 2019). We observed genes with higher average expression variation were more likely to be multi-tissue over-expression outliers (Supplemental Figure 1B).

### Outliers are enriched for rare variants on the X-chromosome

Previous studies have reported enrichment of rare variants near multi-tissue gene expression outliers of autosomal genes (Ferraro et al. 2020; Li et al. 2017). To conduct X-chromosome analyses required additional quality control, and variant filtration (see Methods) (Khramtsova et al. 2023). We subsequently tested for corresponding enrichment of rare variants on the X-chromosome by identifying what proportion of multi-tissue outliers had nearby rare variants compared to the proportion of non-outliers with a nearby rare variant. We observed significant enrichment of gene-proximal rare variants with minor frequencies less than 0.1%, while common variants were not enriched (Figure 1B). Enrichment of rare variants were largely driven by under-expression outliers (Supplemental Figure 2). We observed the point estimate on the X-chromosome to be higher than on the autosomes and chr7, however this was not significant (subsetted to be of equal size to chrX).

We next investigated if outliers were enriched for rare variants within specific annotations (+/− 5kb). For the X-chromosome, we observed that more outliers had a rare variant in either a VEP-annotated transcription factor binding sites, upstream gene, regulatory region, 3’ untranslated region, and 5’ untranslated region as compared to non-outliers (Supplemental Figure 3A). These enrichments were seen on the autosomes, but were not significant on chr7 when considered alone (Supplemental Figure 3B,C).

### Estimated impact of sex-stratification on outlier discovery

Given that females carry two X-chromosomes while males carry just one, we aimed to estimate the impact of sex on outlier discovery. We reasoned that sex stratification could impact outlier discovery in the following ways: a gene could be an outlier in a sex-stratified group but not in the combined group, a gene could be an outlier in a combined group but not in the sex-stratified group, a gene could be an outlier in both the combined and sex-stratified groups but at different strengths, or there could be no substantial impact. First, we simulated how sex-stratification could impact outlier discovery.

To simulate the impact of sex-stratification, we assumed males and females were drawn from a normal distribution, and the combined group was drawn from a mixture model composed of these two aforementioned distributions. We then calculated outliers for *sex-stratified* groups (males or females separately) of equal sample size, and then created a subsetted *sex-combined* group (males and females together) of sample size equal to the sex-stratified groups. We next asked how a shift in mean between the male and female distributions would impact outlier discovery assuming constant variance of 1. The larger the difference in the means between males and females, the more outliers were identified in the sex-stratified groups. Further, we identified outliers in the sex-combined group that were not outliers in the sex-stratified group. We observed this effect up to a mean difference of 1.9 where beyond this point the variance in the sex-combined distribution is large enough that any value called an outlier in the combined group would also be considered an outlier in the sex-stratified group (Supplemental Figure 4A).

We estimated how a shift in variance impacts outlier calling assuming a constant mean for both distributions of 0, and setting the variance of one distribution to be fixed at 1. The change in proportion of outliers due to a difference in variance was a fixed proportion; for example if the variance of the male distribution is 3 and the variance of the female distribution is 6 this was equivalent to if the variances were 1 and 2, respectively (Supplemental Figure 4B). The proportion of outliers lost from the sex-combined group and gained in the sex-stratified group increased as the difference in variance increased, but the proportion of outliers lost happened at a larger rate than the proportion of outliers gained.

### Sex masks gene expression outlier discovery and impacts rare variant enrichments

We tested the impact of sex stratification on our multi-tissue outlier detection using GTEx data. We observed males appeared to carry slightly more outliers than females on the X-chromosome driven by a significantly higher number of under-expression outliers (Fisher’s exact test p=0.01) (Figure 2A; Supplemental Figure 5). On the autosomes there were no significant differences in the number of outliers between males and females. Further, despite equal sample-size, there were fewer outliers in the male (p=0.001) and female (p=6e-05) groups on the autosomes as compared to outliers in the both group, largely driven by significantly fewer over-expression outliers on the autosomes in the female (p=7e-05) and male (p=0.006) groups as compared to the both group. As over-expression outliers are less likely to be driven by rare variants, this indicates stratification can improve detection of genetically-driven outliers.

**Figure 2:**
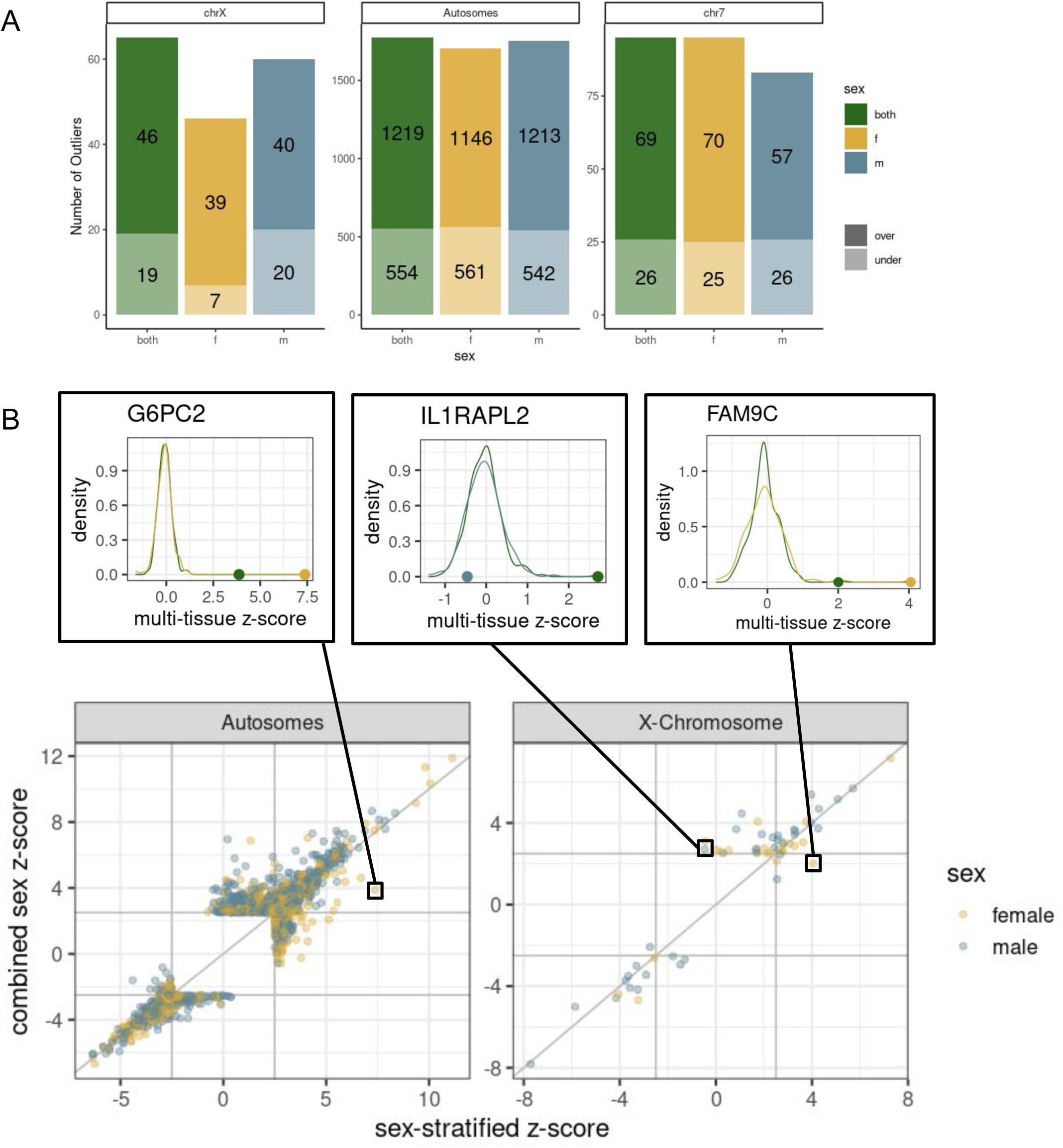
Impact of sex-stratification on outlier detection. a, The number of outliers across the autosomes, chromosome 7, and the x-chromosomes. Under-expression outliers are lighter and on the bottom, over-expression outliers are darker and on the top. b, for an individual-gene pair the z-score in the within-sex group (male or female) as compared to the between-sex group. The individual-gene pair points are colored by their sex. Three examples are highlighted above.

We investigated genes detected as an outlier for an individual in only the sex-stratified or sex-combined group. The vast majority of genes maintain consistent z-scores regardless of group. We observed only 0.1% (27) X-chromosome and 0.2% (951) autosomal genes change outlier status between groups out of all gene-individual pairs that were an outlier in at least one group (Figure 2B; Supplemental Figure 5, Supplemental Table 1). Gene-individual pairs consistently outliers in both the combined-sex and sex-stratified group showed an average z-score shift of 0.4 on the X and autosomes, and were mostly over-expression outliers. For the gene FAM9C, one female is an outlier in the female-only group (z=4.1) but not in the combined group (z=2.0). FAM9C evolved independently on the X after recombination stopped with the Y, and often escapes XCI (Wilson Sayres and Makova 2013). For the gene IL1RAPL2, one male was an outlier (z=2.7) in the combined group but not in the sex-stratified group (z=−0.5); this gene is in a region associated with X-linked nonsyndromic cognitive disability (Valnegri et al. 2011; Zhang et al. 2010). Another gene, G6PC2 is an outlier for the female (z=7.4) and combined (z=3.9) groups. This gene is associated with release of glucose into the bloodstream and has been shown to have sex-specific effects in mice and chicken (Bosma et al. 2020; Wang et al. 2020; Boortz et al. 2017). We observed no genes with an outlier status change in the opposite direction, further supporting that improvements in outlier detection due to sex-stratification are due to uncovering previously masked effects.

Given sex-stratification impacts outlier detection, we investigated its impact on the enrichment of the proportion of outliers with a nearby rare variant compared to non-outliers. Across the autosomes, we observed no significant differences in enrichment (Figure 3). We have previously shown that outliers are more enriched for having nearby deleterious variants (higher CADD score) (Bonder et al. 2021). Males were enriched for rare deleterious (CADD>15) variants on the X-chromosome (Figure 3; Supplemental Figure 6). This enrichment was driven by under-expression outliers (Supplemental Figure 7). The point estimate for males was higher than females (males = 62.4, females = 5.5; Supplemental Figure 6). In the male group, we observed a larger enrichment on the X-chromosome for very rare variants as compared to the autosomes (Supplemental Figure 6).

**Figure 3:**
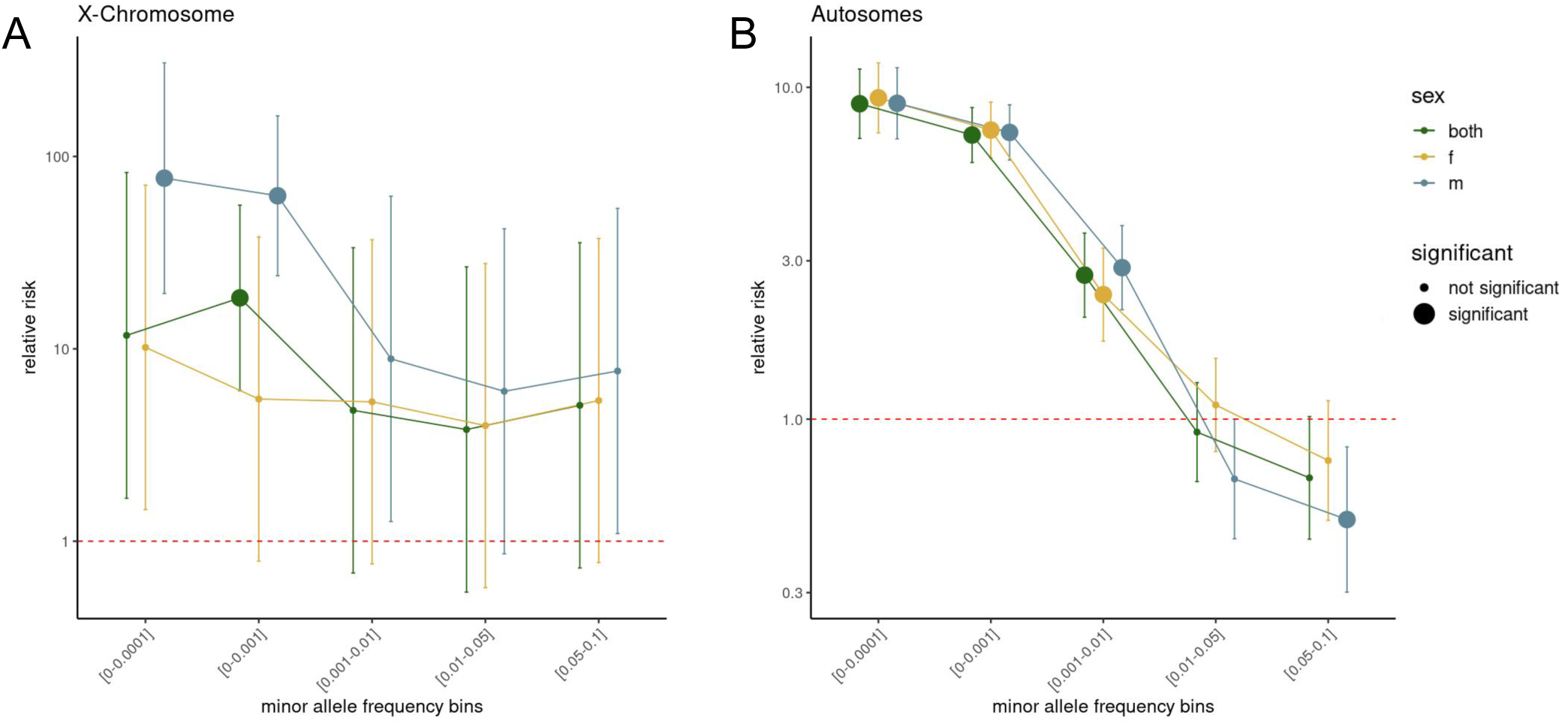
Impact of sex-stratification on relative risks. The relative risk of outliers having a nearby rare variant stratified by sex for **a,** the X-chromosome and **b,** the autosomes across minor allele frequency bins. This is plotted on a log-scale, and the point size indicates statistical significance.

### Discovery of sex-specific functional rare variants

Given the prevalence of sex-biased gene expression in human tissues, we hypothesized that rare variants could also manifest sex-specific effects (Oliva et al. 2020). To test this, we trained a hierarchical Bayesian model, RIVER, using multi-tissue expression data from both sexes. RIVER significantly outperformed genomic annotation models (GAM) in predicting outlier status of individuals with shared rare variants (Supplemental Figure 8) (Li et al. 2017). We then applied this model to estimate the posterior probability for variants driving outlier status in female and male individuals separately. Overall, we scored 1.9 million rare variants on autosomes and 29 thousand variants on chromosome X. Sex-biased variants were defined to be those with a difference in the posterior probability more than 0.2.

The vast majority of variants shared similar functional predictions in both sexes (99.5% with posterior probability difference < 0.1); however, we identified 5524 variants with sex biased posteriors. For increased confidence, we then only included variants that were sex-biased in at least one of five permutations of the sex labels, reducing our set to 753 variants across 464 genes, none of which included the X-chromosome (Supplemental Table 2, Figure 4a, see Methods). There were a similar number of female-biased and male-biased functional rare variants (Supplemental Table 3). There was no ontological enrichment for genes with a sex-biased variant (Yu et al. 2012).

To investigate the functional impact of sex-specific rare variants, we curated a comprehensive set of annotations. We then examined the proportion of functional rare variants (defined as RIVER posterior > 0.2) with a given annotation (missense, intronic, etc) and did so for those that were female-biased, male-biased, and not sex-biased. Here, we defined the background as all scored variants (appearing in both females and males), and sex-shared variants as those with large posteriors (> 0.2) in both sexes. Several variant types were enriched for being sex-biased and functional as compared to non-functional, such as transcription factor binding sites, stop-gained, and splice sites (Figure 4b, Supplemental Figure 9a,b). Non-functional variants were enriched in non-coding and intergenic regions (Figure 4b, Supplemental Figure 9a,b).

**Figure 4:**
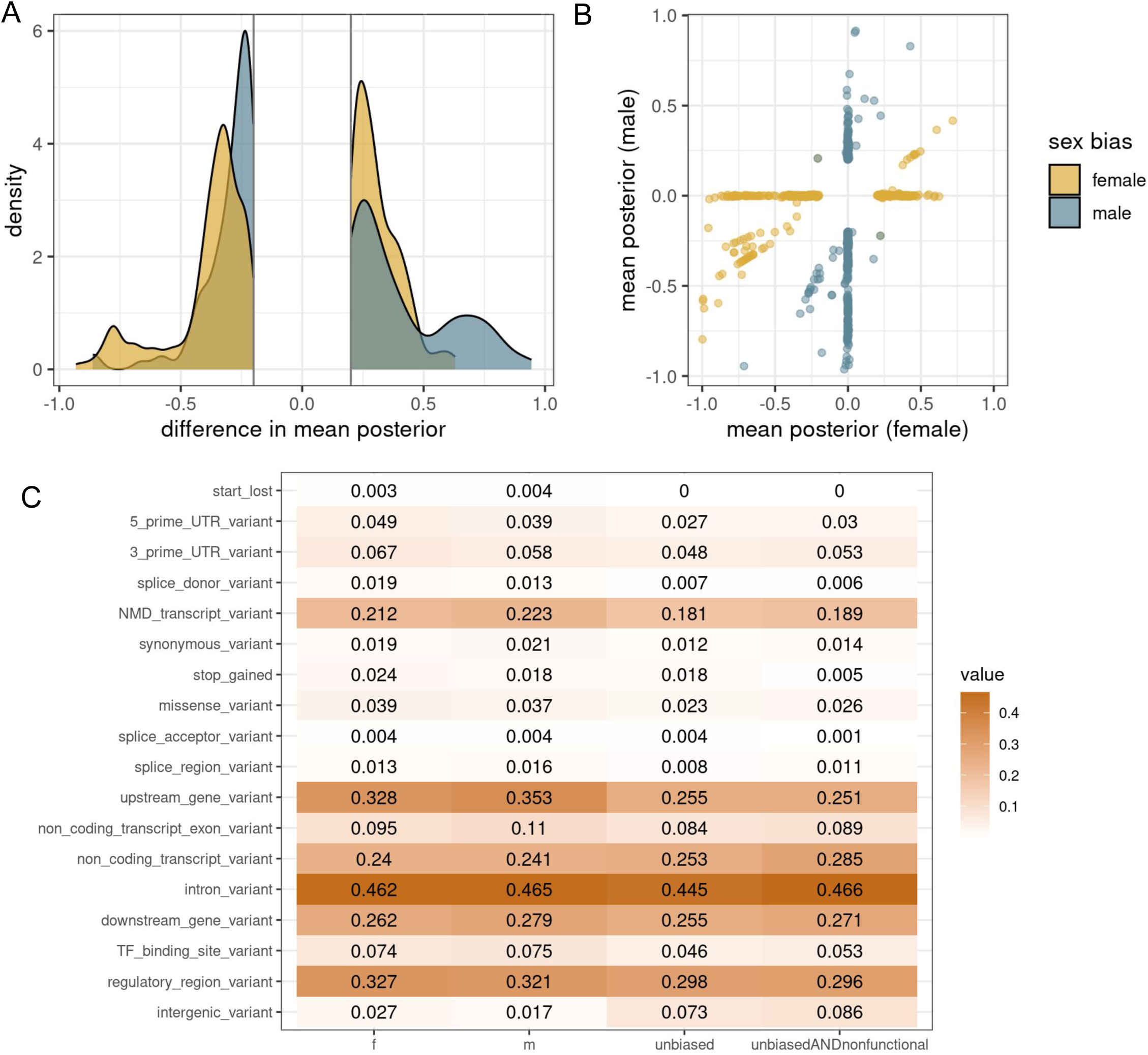
Sex-biased functional rare variants. For predicted functional (abs(beta)>0.2) sex-biased (absolute mean difference > 0.2), the (**a**) distribution of difference in mean posteriors between males and females, and (**b**) the values of the posteriors in each sex. **c**, For each given variant category, the proportion of rare variants that had this annotation. Note that variants can be in multiple categories. This is calculated for female-biased, male-biased, unbiased, and unbiased and also nonfunctional rare variants.

There was a significant overlap of 11 sex-biased eQTLs genes from Oliva et al. and our sex-biased rare variant list (hypergeometric test p=0.008), and a significant overlap of 37 sex-biased eQTL genes from Jones et al. (hypergeometric test p=5.7e-6; Supplemental Figure 10a) (Jones et al. 2024; Oliva et al. 2020). The gene TDRD12, which has been linked to fertility, overlapped in all three sets (Babakhanzadeh et al. 2020; Dai et al. 2017). When considering tissue-specific scores from Oliva et al. the adrenal gland, which has high hormonal activity and sexual dimorphism, had the largest correlation to the sex-biased rare variants (Supplemental Figure 10b, Supplemental Table 4) (Lyraki and Schedl 2021).

We identified two rare variants in two genes that were predicted to exhibit sexual antagonism (absolute mean posterior greater than 0.2 in both sexes with different directions). The variant chr11:75430623 was predicted to increase expression of KLHL35 in females and decrease expression in males. Increased expression of KLHL35 has been linked to survival levels for lung adenocarcinoma with a TP53 mutation (Fu et al. 2021), and females have been shown to have better survival of this disease (Freudenstein et al. 2020). The stop-gain variant chr1:171636338 was predicted to increase expression in females and decrease expression in males for the gene MYOC, and has a CADD score of 37 and is likely-pathogenic for glaucoma. Mutations in this gene can cause glaucoma, and variants in other glaucoma-causal genes have been linked to sex-bias of this disorder (Suri et al. 2009). MYOC has also been linked to sex-determination in zebrafish (Atienzar-Aroca et al. 2021).

### Sex-specific functional rare variants occur in genes with known sex-biased drug interactions

Rare variants account for an estimated 30-63% of variation in drug response (Kozyra et al. 2017; Al-Mahayri et al. 2020; Bertholim-Nasciben et al. 2023), and approximately half of known pharmacogenes are exclusively driven by rare variants (Ingelman-Sundberg et al. 2018). It has further been observed that 77-96% individuals contain at least one deleterious rare variant in an actionable pharmacogenetic gene (Yu et al. 2021; Abouelhoda et al. 2024; Nunez-Torres et al. 2023; Chan et al. 2022). We sought to determine if the role of predicted functional sex-biased variants were relevant to pharmacogenetics. To do so, we compared 734 functional sex-biased rare variants (from 449 genes) to 4953 functional non-sex-biased rare variants (from 2246 genes) across pharmacogenetic databases.

To connect functional rare variants to genes that are of pharmacogenetic interest we used the Drug Gene Interaction database (DGIdb), which contains 13,186 drugs with known 6,888 gene interactions (Cannon et al. 2024). We identified 105 sex-biased functional rare variants from 55 unique genes that interact with 570 unique drugs (Figure 5a, Supplemental Figure 11a, Supplemental Table 5). One such variant was the intronic variant chr16:28607045 which was predicted to increase expression of SULT1A1 only in females. This gene plays a known role in estrogen metabolism (Spink et al. 2000; Liu et al. 2017). Of these unique drugs, 491 interacted with a gene that had a female-biased functional rare variant while just 176 interacted with a gene that had a male-biased functional rare variant (Supplemental Figure 11a, Supplemental Table 5).

**Figure 5:**
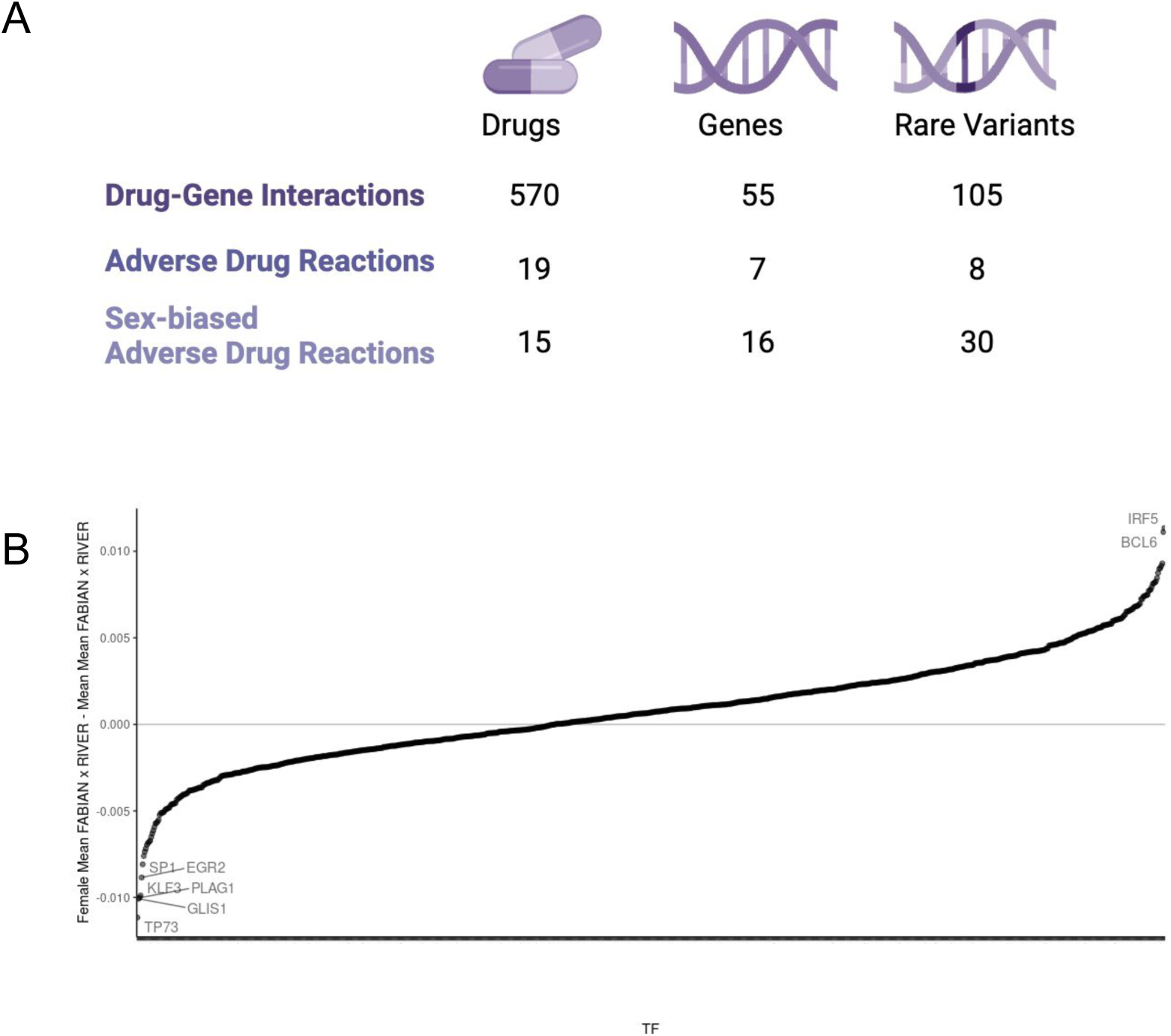
Sex-biased functional rare variants. **a,** Across the Drug Gene Interaction Database (DGIdb), the FDA list adverse drug reactions, and adverse drug reactions with known sex-biases from Zucker and Prendergast 2020, the number of drugs and genes that were linked to a sex-biased rare variant. **b,** For FABIAN scores x RIVER scores, the median gene-summarized difference between sexes. This is ordered with positive values representing increased binding in females as compared to males, and negative being the reverse.

It has been observed that adverse drug reactions occur at a higher rate in women (Moyer et al. 2019), and there are over 20,000 predicted adverse drug effects with sex differences (Chandak and Tatonetti 2020). The FDA lists 131 drugs with adverse drug reactions that have been linked to a gene (Center for Drug Evaluation and Research 2024). We identified 8 functional sex-biased rare variants from 7 genes linked to 19 drugs with adverse drug reactions (Figure 5a). Despite having a similar number of functional sex-biased rare variants (5 female and 3 male), females had more associated drugs with these adverse reactions (17 female, 2 male) (Supplemental Figure 11a, Supplemental Table 5). Of 59 drugs previously identified to have sex-differences in adverse drug reactions, 15 were linked to a gene which also had a sex-biased functional rare variant (Figure 5a, Supplemental Figure 11a, Supplemental Table 5) (Zucker and Prendergast 2020). This was enriched as compared to drugs linked to genes with non-biased functional rare variants (relative risk = 1.67, p-value = 0.005) (Supplemental Table 6).

We identified four functional rare variants in the gene CBR1, all of which were male-biased and predicted to reduce expression (Supplemental Figure 11b). CBR1 is a reductase of doxorubicin and upregulation of this gene drives doxorubicin resistance (Matsunaga et al. 2015) which leads to increased myotoxicity, of which there are known sex differences for this drug (Montalvo et al. 2021). Therefore reduced expression of this gene is preferred, and reduction of CBR1 in mice improved doxorubicin efficacy only in male mice (Freeland 2011).

### Identification of sex-specific TF networks

Given that transcription factor (TF) activity has been linked to sex-biased gene expression across human tissues, we hypothesized that sex-biased variants in regulatory regions may inform discovery of TFs and TF networks with sex biases (Lopes-Ramos et al. 2020). To this end, we first performed a motif enrichment analysis using HOMER with 100 bp window on both sides of female- and male-biased rare variants (Heinz et al. 2010). The top known motif was a target of the gene SIX1 (Supplementary Figure 12a). SIX1 is modulated via a microRNA intermediated by the highly sex-biased gene XIST, SIX1 can modulate a key regulator of the sex determination pathway SRY, and SIX1 knockout reduced male gonadal development in mice (Liu et al. 2020; Fujimoto et al. 2013). The top *de novo* enriched motif was a target of HAND2 (Supplementary Figure 12b). This gene plays a large role in pregnancy and the menstrual cycle, is a downstream target of estrogen and progesterone signaling, and can also regulate estrogen (Oh et al. 2023; Marinić et al. 2021).

We then use FABIAN-variant, an in-silico modeling approach based on a large collection of TF motif databases that identifies TFs likely disrupted by variants, to further understand the impact of the sex-biased rare variants on TFs (Steinhaus et al. 2022). We multiplied the FABIAN-variant score with the RIVER score, and signed it such that positive meant potential gain of binding and negative meant potential loss of binding. With this, we were able to identify TFs for target genes whose motifs were predicted to be disrupted by sex-biased variants (Supplemental Figure 13a). For example, YY1 – a TF known to regulate sex-biased transcription – was identified as a TF targeting a gene with predicted sex-biased rare variants (Supplemental Figure 13a) (Chen et al. 2016). We further identified TFs with a difference in gain or loss of binding for sex-biased rare variants (Supplemental Table 7). For example, BCL6 was identified as predicted increased binding in females but decreased binding in males (Figure 5a, Supplemental Figure 13b, Supplemental Table 7). BCL6 has been shown to preferentially bind to female-biased targets (Zhang et al. 2012). SP1, which has preferential male binding, is known to bind SRY (Desclozeaux et al. 1998). Overall, our data suggest that rare variants can mediate sex-specific regulatory functions through sex-biased transcription factors.

## Discussion

Previous studies have examined the impact of rare variants across the autosomes using outlier-based approaches (Smail et al. 2022; Frésard et al. 2019; Brechtmann et al. 2018; Ferraro et al. 2020; Li et al. 2023). However, to date, none of these approaches have been applied to the X-chromosome. In this work, we extended this approach to identify multi-tissue outliers on the X-chromosome, and found these outliers were enriched for nearby rare variants.

We demonstrated that rare variants are more enriched for gene expression outliers on the X-chromosome in males as compared to females. This follows the intuition that the additional copy of the X-chromosome in females may compensate for some extreme variant effects, and follows patterns of X-linked mutations having stronger impacts in males (Arnold et al. 2016). We also observed significantly fewer over-expression outliers on the autosomes in the sex-stratified groups as compared to the sex-combined group, indicating that this approach removes some false outliers. This emphasizes the importance of X-chromosome and sex-stratified analyses particularly for genes with large differences in mean or variance between sexes.

Sex biased eQTL studies have demonstrated the power challenges with uncovering genetic effects. However, we reasoned that machine learning approaches that combined molecular outliers and variant annotation, when trained on sex-stratified datasets, may elucidate candidate causal variants. Using this approach, we identified 753 rare variants with different predicted effects between males and females, and observed that these sex-biased rare variants are enriched in specific regulatory regions and annotations. We identified just two genes, KLHL35 and MYOC, that had rare variants with predicted effects in opposite directions between males and females, supporting the hypothesis by Zhu et al. that genetic variants modify sex differences primarily through amplification (Zhu et al. 2022). As machine learning models predicting variant effects evolve, we highlight the advantages to considering sex as an important variable, not just to regress out, but to stratify in training and testing. For example, in an Alzheimer’s dataset, sex-stratified training led to better disease prediction and highlighted different underlying pathways (Bourquard et al. 2023). This indicates that sex-stratified model training can elucidate sex-specific variant effects, and demonstrate a strong impact for X-chromosome variants.

Biological sex has been documented to influence pharmacokinetics, pharmacodynamics, adverse drug reactions, and toxicity, but there remain challenges identifying the biological mechanisms (Ruzzo et al. 2019; Gaignebet and Kararigas 2017; Hampel et al. 2018). Rare variants are also known to have a large impact on pharmacogenes (Ingelman-Sundberg et al. 2018). We identified a functional sex-biased rare variant in 0.8% of known pharmacogenes in DGIdb (Cannon et al. 2024). Genes with adverse drug reactions were enriched for containing a sex-biased functional rare variants, including in clinically actionable genes. This suggests functional rare variants that act differently between males and females may help to explain sex-differences in pharmacogenetics and adverse drug reactions. However, such candidates remain to be functionally evaluated.

One hypothesis for how sex-biased variants ultimately impact expression is through sex-biased expression of transcription factors. Jones et al. found that sex-biased eQTLs are predominantly targeted by transcription factors with sex differences (Jones et al. 2024). We found that sex-biased rare variants significantly overlapped with sex-biased common variants in their target genes. In this study, we were able to replicate sex-biased transcription factors using enrichment analysis of genomic regions defined by sex-specific rare variants. These results provide intriguing possible mechanisms of sex-biased transcription factor regulatory networks across human tissues (Lopes-Ramos et al. 2020), in which both common and rare genetic variants contribute to sex-biased transcription profiles through modulation of transcription factor binding in sex-specific manners (Jones et al. 2024). Importantly, permutation of sex labels failed to replicate these findings, suggesting that our sex-stratified RIVER models are robust in discovering sex-specific gene regulatory networks anchored by rare genetic variants. Screening for motif enrichment around sex-biased variants identified several targets of genes involved in highly sex-biased genes, and relevant transcription factors with known sex-differences in binding, such as BCL6. Future analysis on the differential effects of sex-specific rare variants on complex traits could further elucidate the specific TFs and relevant tissues underlying observed sex differences.

There are several limitations to consider. Our analyses refer to XY males and XX females, and excludes intersex individuals or individuals with other combinations of sex chromosomes due to a lack of power and as such does not capture the full spectrum of individuals. A further limitation is the GTEx population is primarily European, and we have filtered to only Europeans given underlying gaps in reference data that would misclassify rare variants. Understanding rare variants in a diverse dataset is both a scientific and ethical priority (Ju et al. 2022; Altshuler et al. 2010). While sex differences are important to understand, as many diseases vary in risk and severity by sex (Khramtsova et al. 2019), it is also important to not over-report sex differences (Garcia-Sifuentes and Maney 2021). Overall, males and females were very similar in our analyses; reflecting Lewontin’s observations for ancestry, when it comes to genetic effects, there are more within-sex differences than between-sex differences (Lewontin 1972; Oliva et al. 2020).

Genetics studies of molecular and trait phenotypes are increasingly elucidating the impacts of low-frequency and rare variants (Wang et al. 2021). However, for the foreseeable future many rare and ultra-rare variants will be difficult to interpret using conventional association testing. In this study, we survey the impact of rare variants on gene expression on the X-chromosome, note how they differ by sex, and implicate further pharmacogenetic and transcriptional factor mechanisms that can create sex-differences.

## Data and code availability

Scripts used for analysis are available at github.com/raungar/rarevariant_x_sex. Data is available to authorized users through dbGaP (phs000424.v8) and on the GTEx portal (https://gtexportal.org/).

## Acknowledgments

Thank you to the donors and family of donors from the GTEx study for making this research possible. This work utilized computing resources provided by the Stanford Genetics Bioinformatics Service Center, supported by NIH Instrumentation Grant S10 OD025082, and would not have been possible without the support of the Stanford SCG cluster system administrators. We thank the Montgomery lab members and Stanford Genetics department members for comments and suggestions throughout the research process. Thank you to R.A.U.’s committee members Noah Rosenberg, Marcia Stefanick, and Jesse Engreitz for feedback over the years. R.A.U. was funded by the Stanford Genome Training Project (T32HG000044) and GREGoR Consortium (U01HG011762). The content is solely the responsibility of the author and does not necessarily represent the official views of the National Institutes of Health.

## Author Contributions

R.A.U. and S.B.M. conceived the study. R.A.U., T.L., and S.B.M. significantly contributed to study design, with feedback from A.B and N.E.. Analyses were performed by R.A.U., T.L., and with mentorship from S.B.M. and A.B. Figures were generated by R.A.U. and the manuscript was primarily written by R.A.U. with major contributions from T.L. and edits provided by S.B.M. All authors provided feedback on the manuscript to improve it.

