## Supplemental Figures for "Functional impact of rare variants and sex across the X-chromosome and autosomes"

**A**

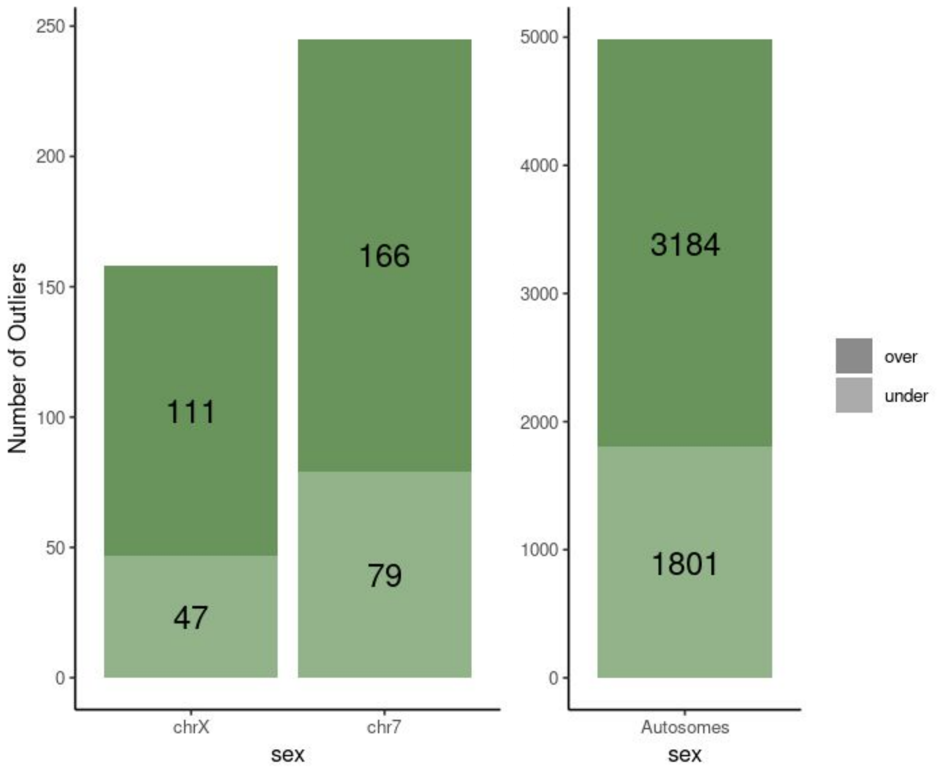


**B**

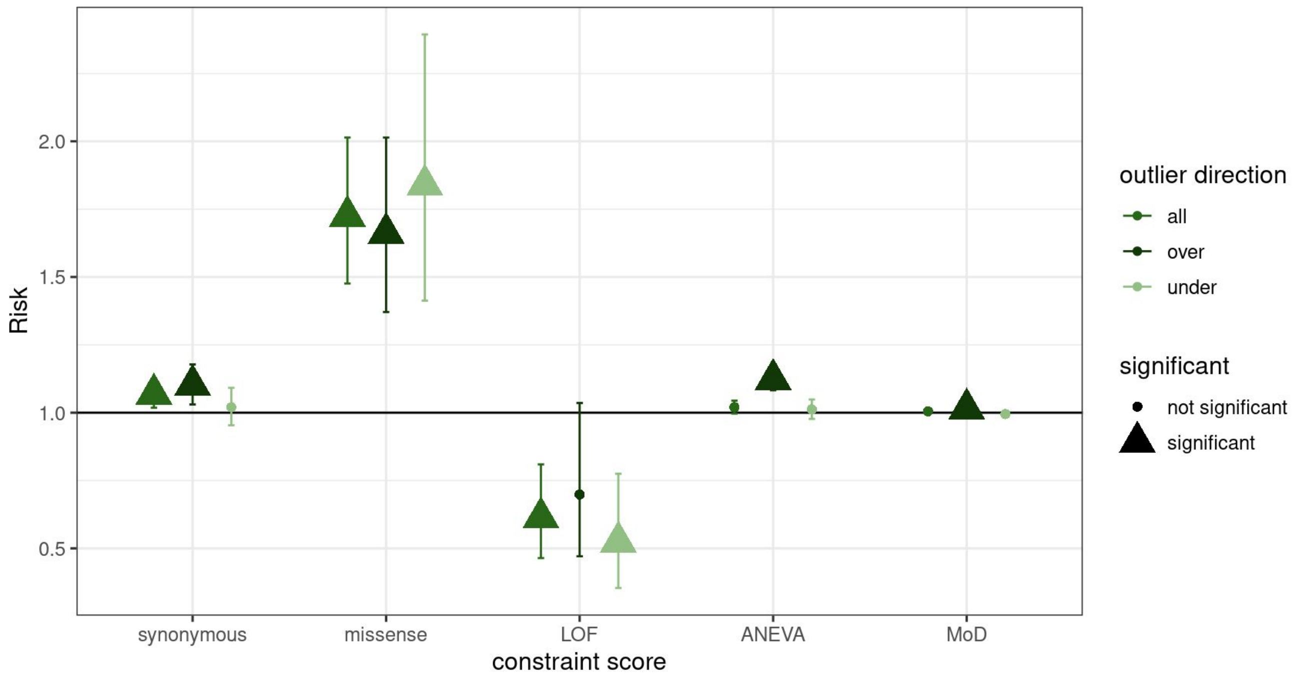


**Supplemental Figure 1: Over and under-expression outlier summary. a,** Number of outliers split by over/under where the lighter shade is under-expression outliers, and the darker shade is over-expression outliers. **b,** relative risk of outliers having a high (synonymous/missense/LOF from Samocha et al. z-score >2, ANEVA from Mohamaddi et al >0.95, MoD from Dong et al > .95) constraint score as compared to a low score (synonymous/missense/LOF from Samocha et al. z-score < -2, ANEVA from Mohammadi et al <.05, MoD from Dong et al <.05)). Significance is Benjamini-Hochberg corrected.

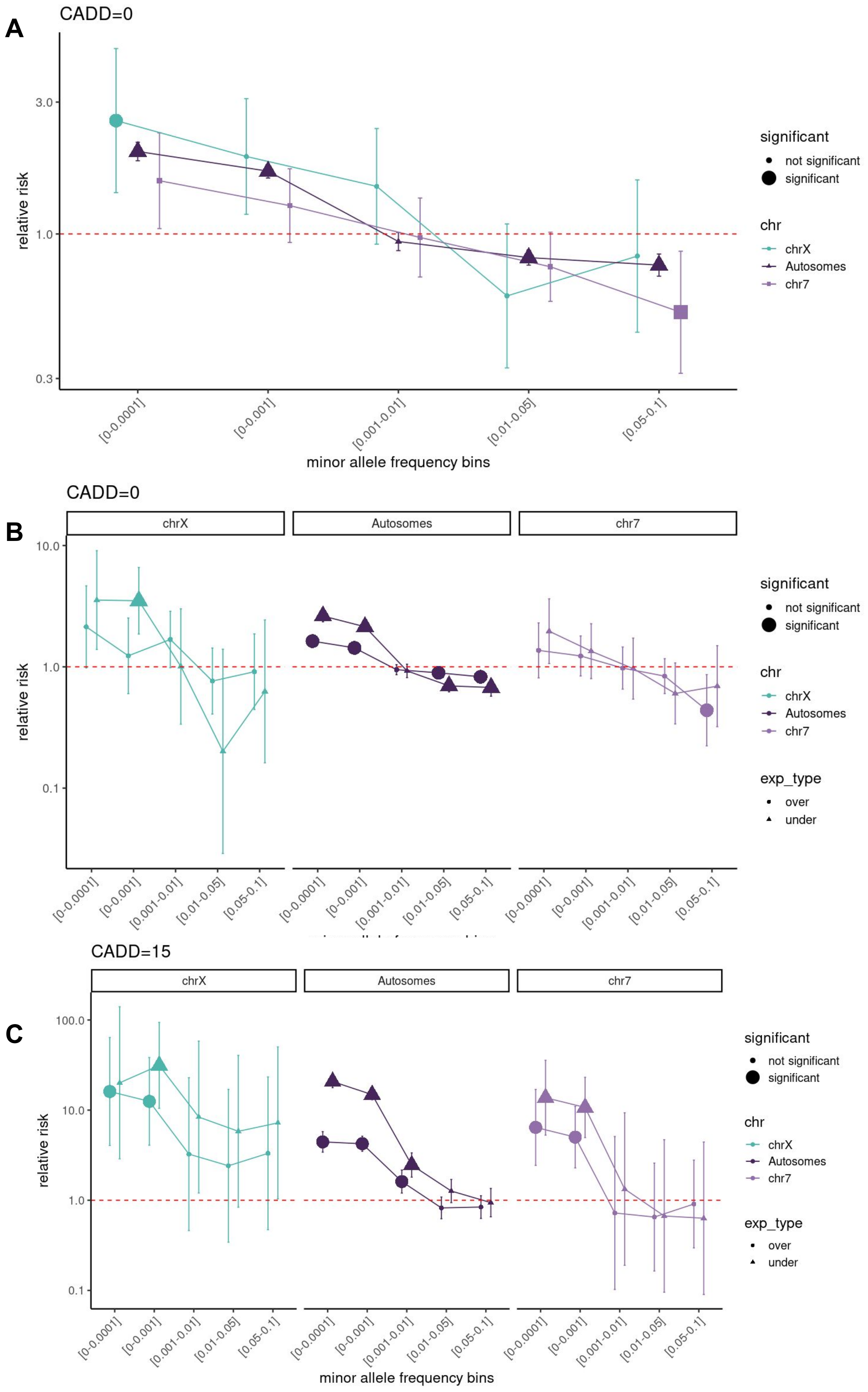


**Supplemental Figure 2: Relative risk enrichments.** These plots show the relative risk of having an outlier with a nearby rare variant as compared to a non-outlier. In a, this is done with no CADD threshold. Further, these enrichments are calculated where the outliers are split to represent either under-expression or over-expression outliers, across the X-chromosome, chromosome 7, and the autosomes at **b,** no CADD threshold and **c,** a CADD threshold of 15.

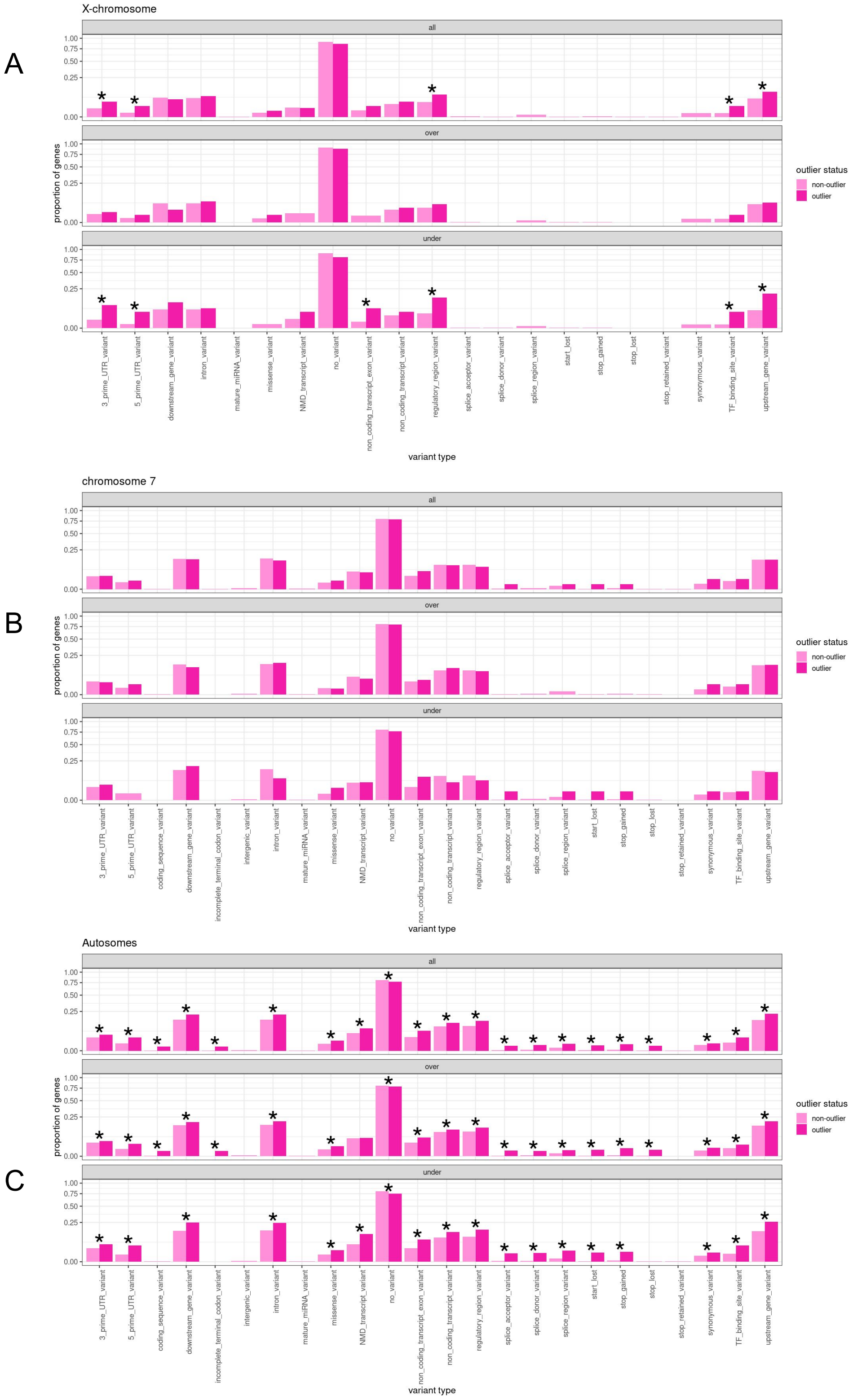


**Supplemental Figure 3: Enrichment of variant types.** The proportion of outliers with a given variant as compared to the non-outliers with a given variant across **a,** the X-chromosome, **b,** chromosome 7**,** and **c,** the autosomes. Significance level is determined by Fisher's exact test and adjusted using the Benjamini-Hochberg approach for multiple testing correction.

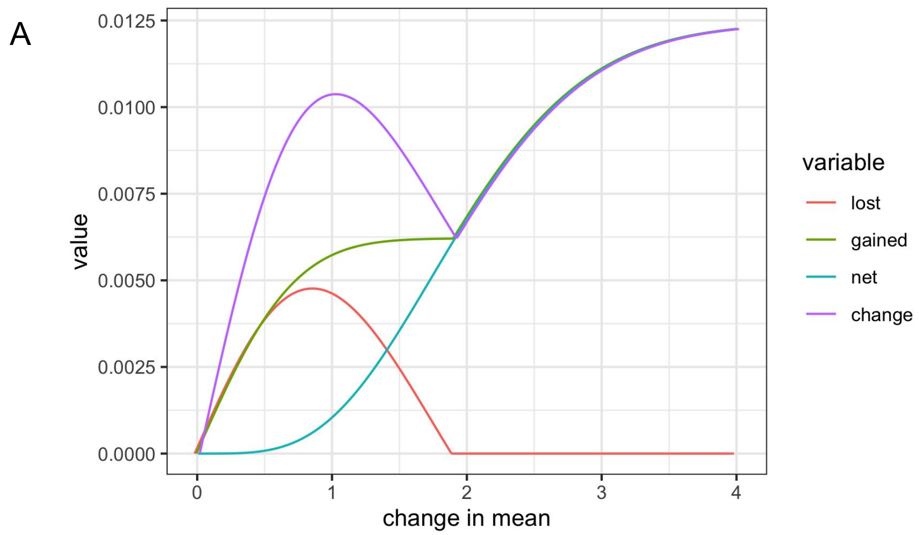


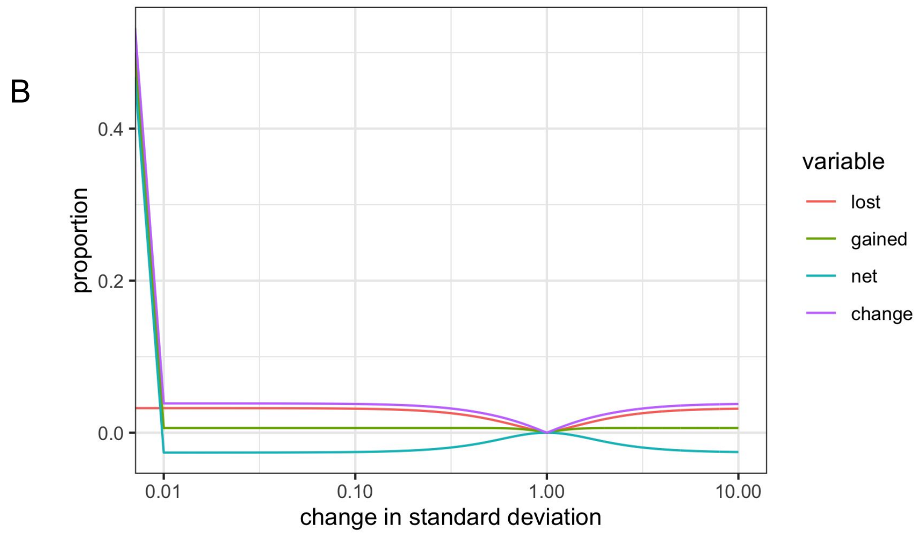


**Supplemental Figure 4: Theoretical impact of sex-stratification.** The impact of changing the **a,** mean or **b,** standard deviation on the proportion of genes that will either lose outlier status and gain outlier status. The change represents the sum of the positive and negative change, while the net change represents the sum of the absolute value of positive and absolute value of negative change.

A

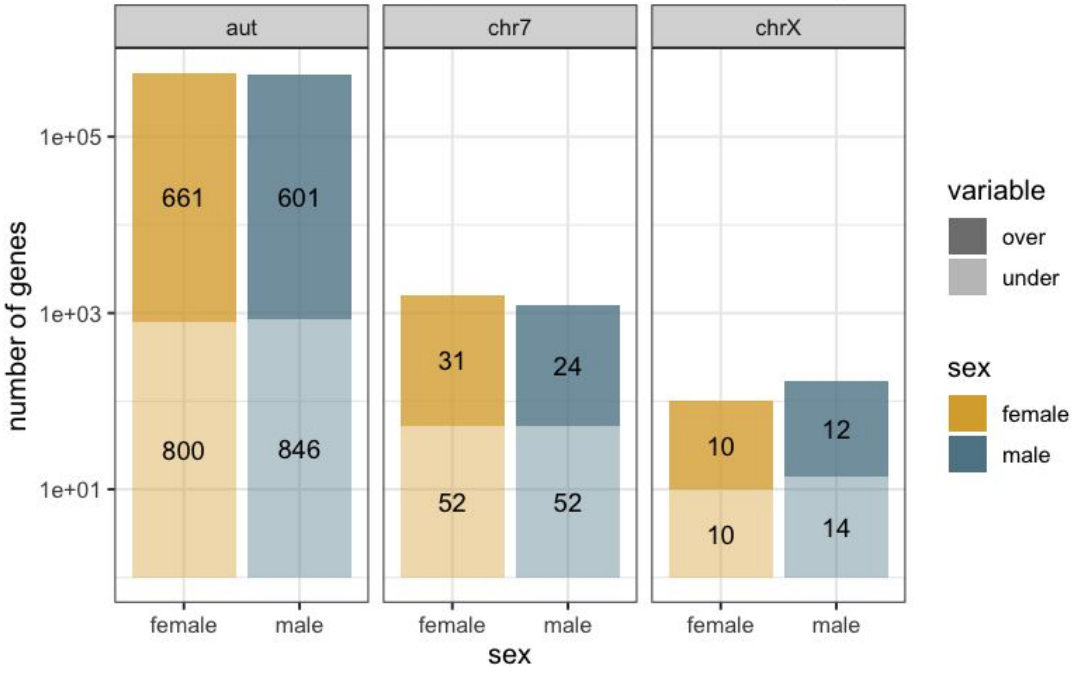


B

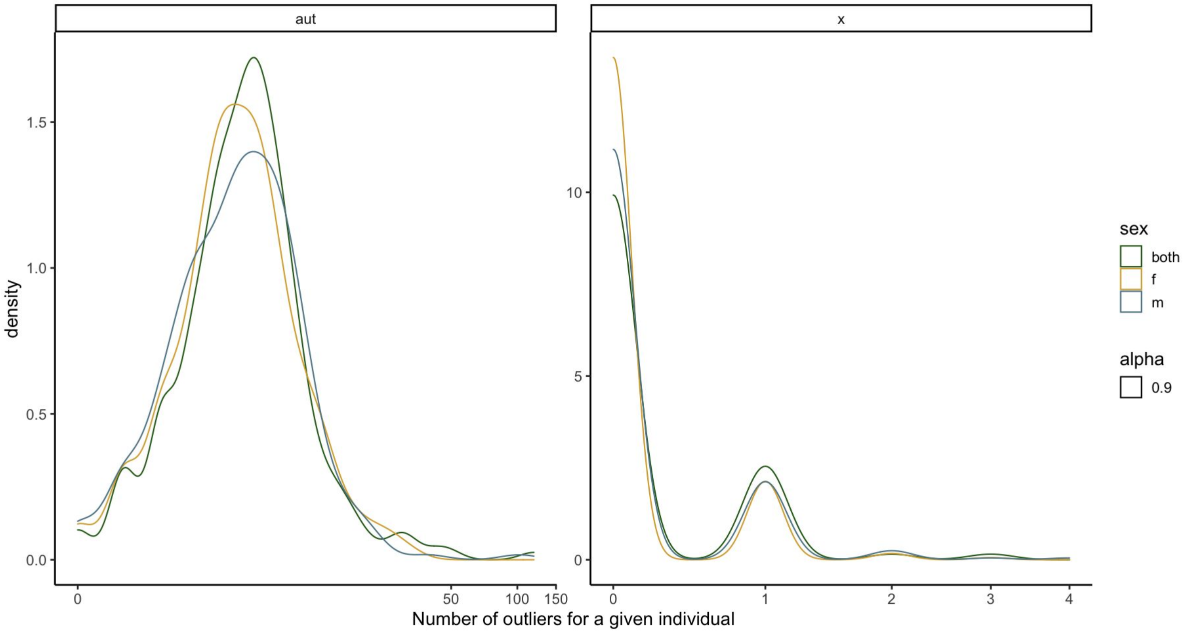


**Supplemental Figure 5: Number of outliers sex-stratified. a,** the number of outliers split by over (darker, on top) and under (lighter, on bottom), across the autosomes, X-chromosome, and chromosome 7. **b,** the distribution of the number of outliers per individual stratified by sex for the autosomes and X-chromosome.

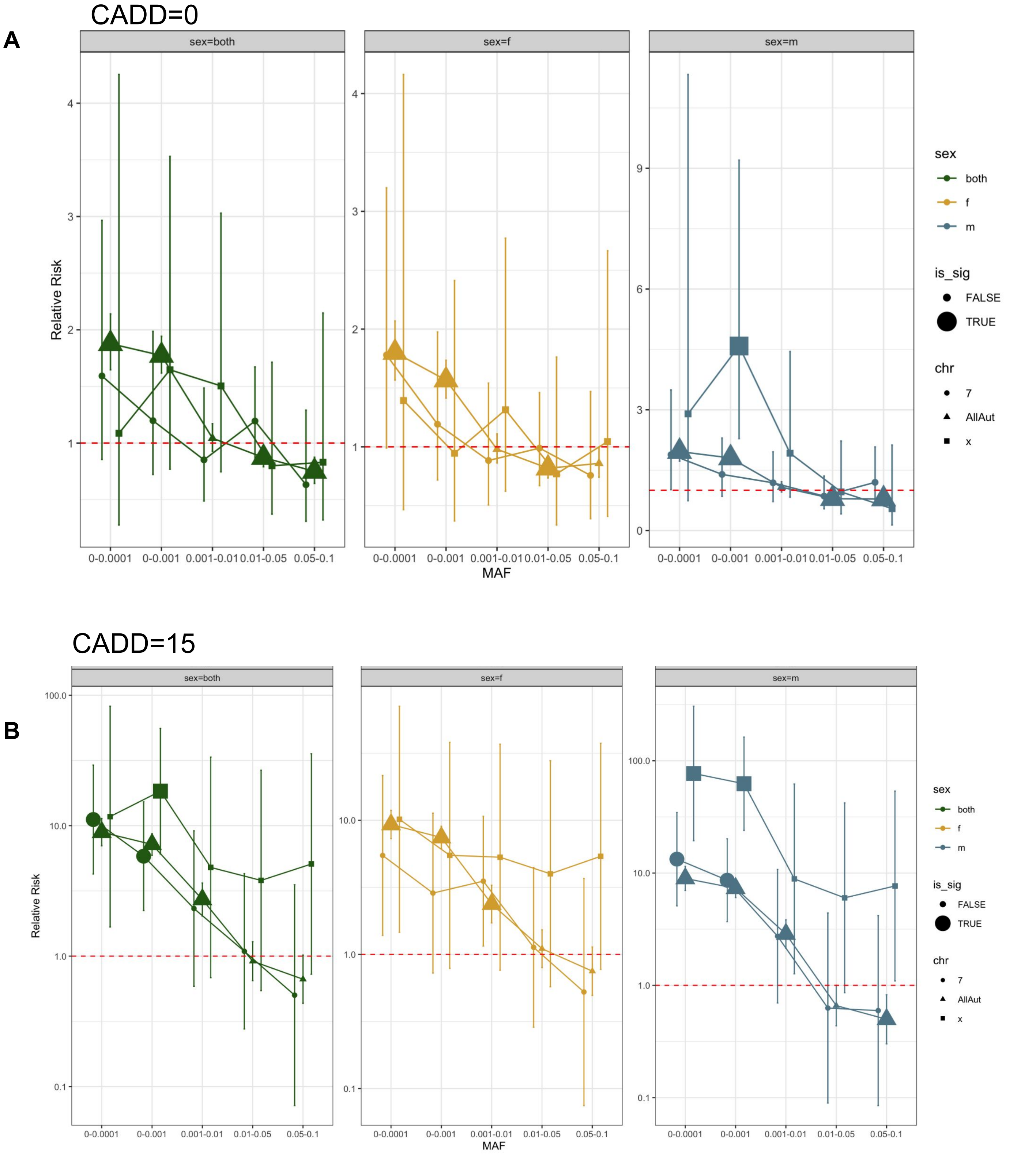


**Supplemental Figure 6: Sex-stratified enrichment scores.** Enrichment for outliers having nearby rare variants as compared to non-outliers at **a,** no CADD threshold and **b,** a CADD threshold of at least 15. This is across the X-chromosome, chromosome 7, and autosomes and stratified by sex. Significance is size of dot.

A

B

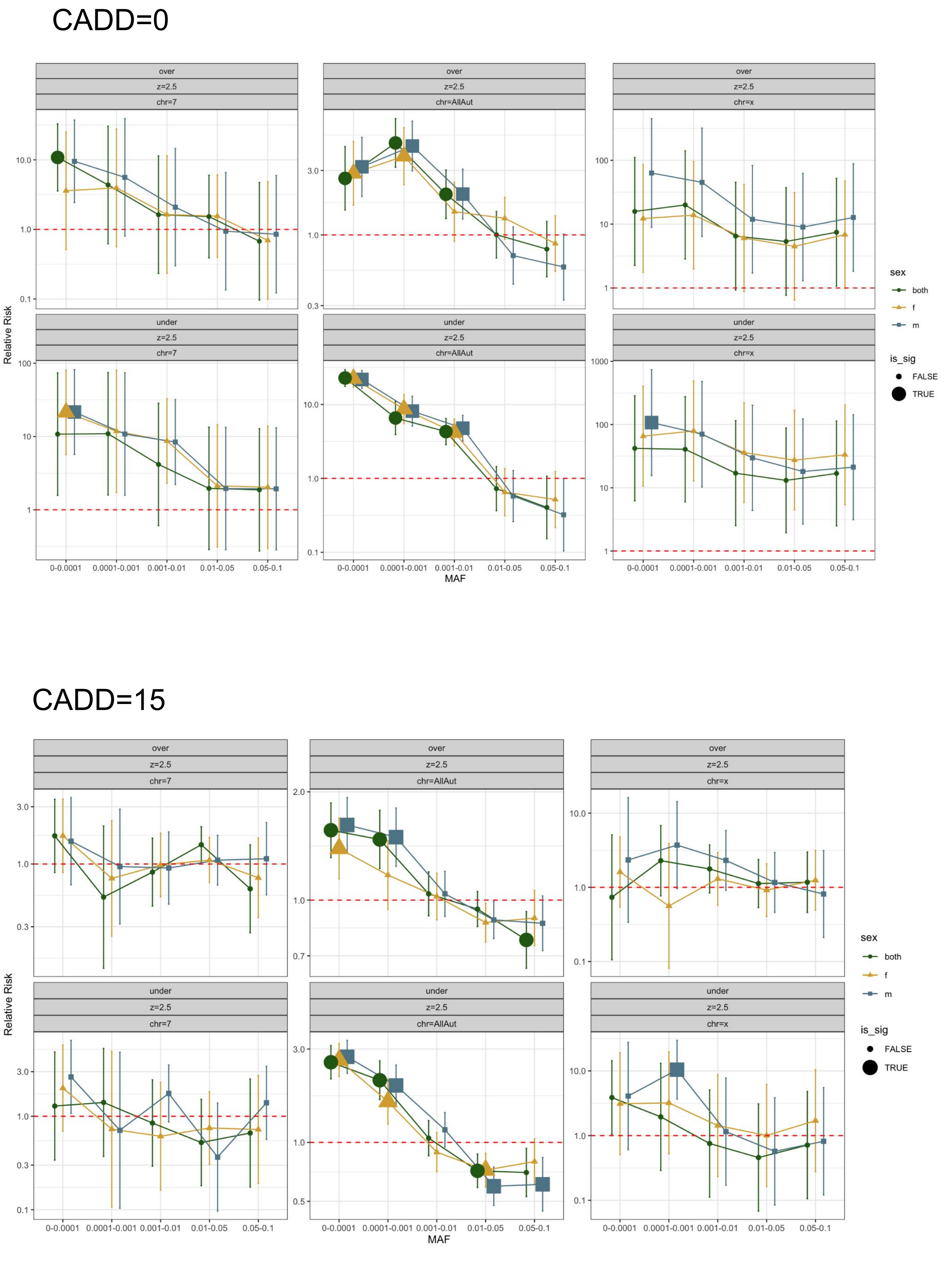


**Supplemental Figure 7: Sex-stratified enrichment scores split by under- and over-expression.** Enrichment for outliers having nearby rare variants as compared to non-outliers at **a,** no CADD threshold and **b,** a CADD threshold of at least 15. This is across the X-chromosome, chromosome 7, and autosomes and stratified by sex. This is for both over- and under-expression outliers. Significance is size of dot.

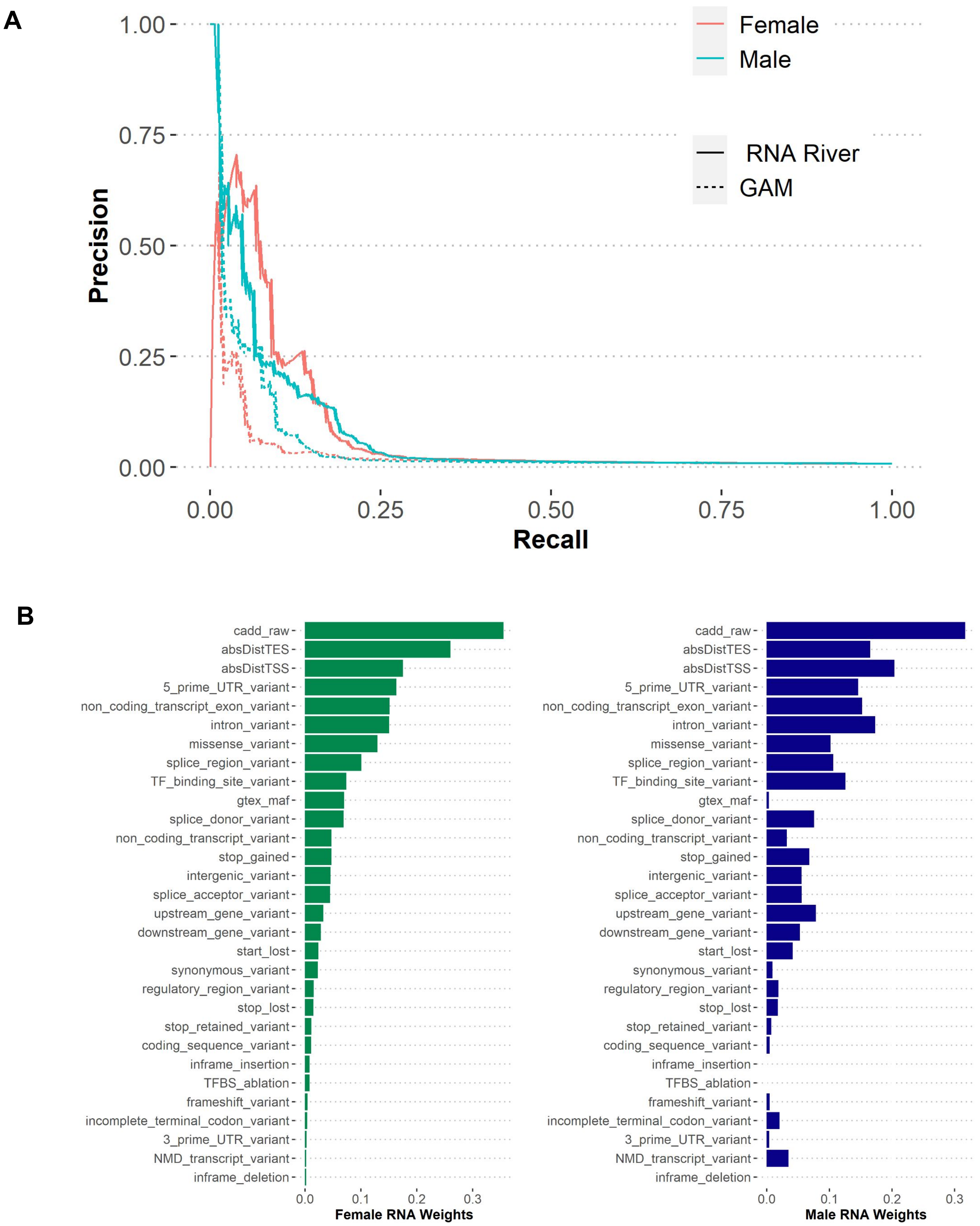


**Supplemental Figure 8: RIVER and GAM modeling.** a) precision-recall curve of female (red) and male (blue) specific RIVER and GAM models. B) Feature weights of genomic annotations learned from sex-specific RIVER models, ranked by female-specific weights.

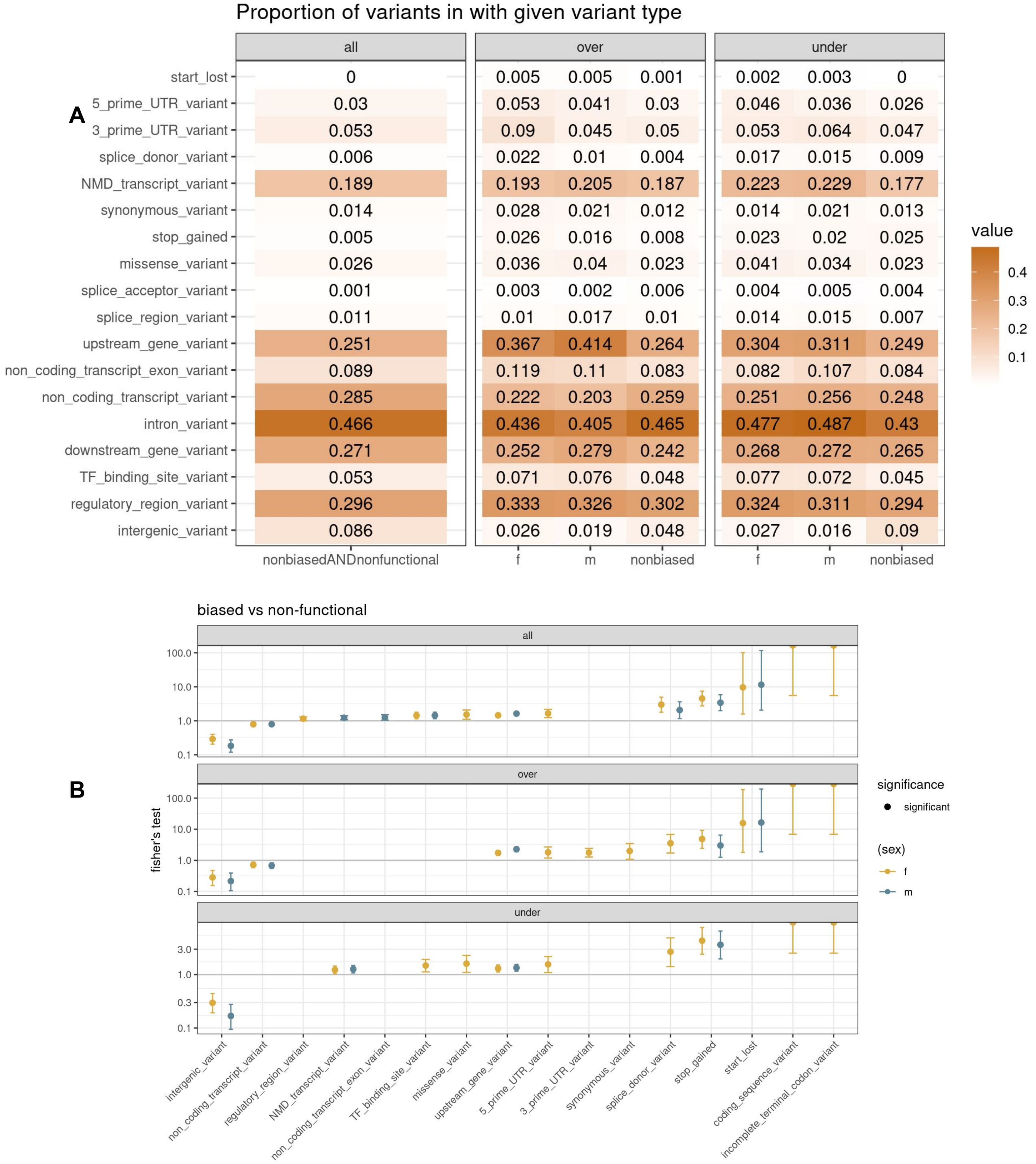


**Supplemental Figure 9:** **a,** For each given variant type, the proportion of rare variants that had this annotation (note: variants can have multiple annotations). This is split by over and under expression across female-biased, male-biased, and nonbiased functional rare variants. **b,** Fisher's test for a sex-biased rare variant being in a particular variant category as compared to a non-functional rare variant. This is done for female-biased and male-biased variants, as well as split for predicted direction of effect (over and under).

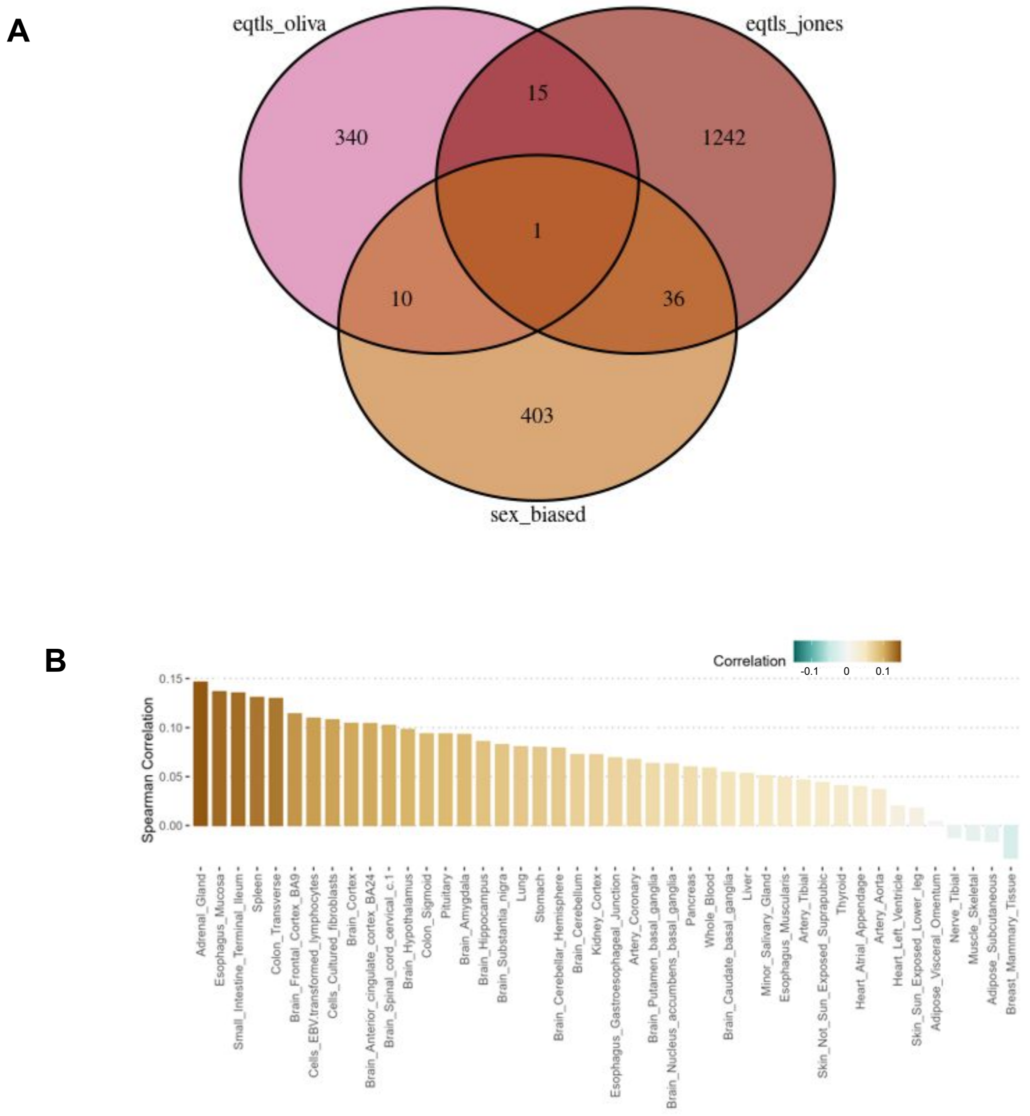


**Supplemental Figure 10:** **a**, Overlap of genes that have at least one sex-biased rare variant, and genes from Oliva et al 2020 with at least one sex-biased eQTL. **b,** Spearman correlation of tissue-specific sex-biased gene scores from Oliva et al 2020 with sex-biased rare variants scores.

**A**

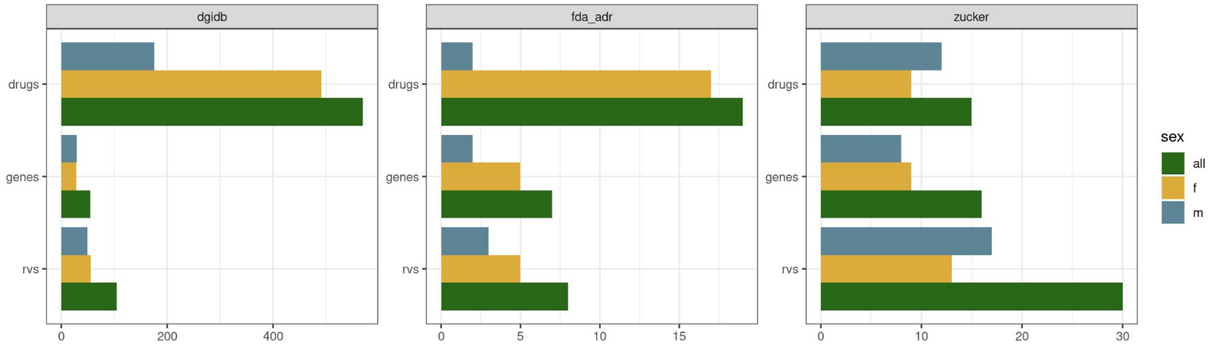


**B** **Screenshot from UCSC Genome Browser**

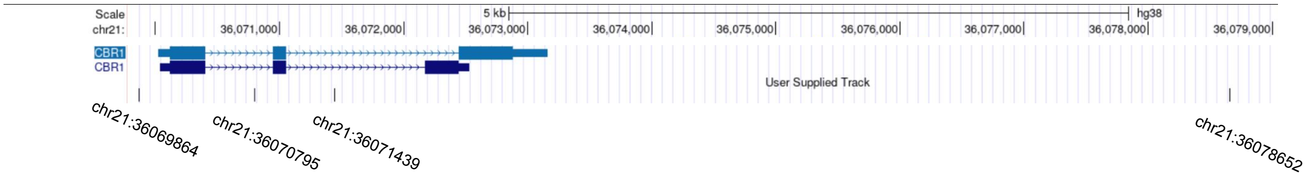


**Supplemental Figure 11: a**, The number of unique rare variants, genes, and drugs that the functional sex-biased rare variants overlapped with in the DGIdb, FDA adverse drug reaction databased, and from Zucker et. al. **b,** Location of sex-biased functional rare variants for the gene CBR1.

**A**

| Rank | Motif | Name | P-value | log P-pvalue | q-value (Benjamini) | # Target Sequences with Motif | % of Targets Sequences with Motif | # Background Sequences with Motif | % of Background Sequences with Motif | Motif File | SVG |
|---|---|---|---|---|---|---|---|---|---|---|---|
| 1 | | Six1(Homeobox)/Myoblast-Six1-ChIP-Chip(GSE20150)/Homer | 1e-4 | -9.450e+00 | 0.0792 | 5.0 | 11.90% | 510.1 | 1.05% | motif file (matrix) | svg |
| 2 | | EKLF(Zf)/Erythrocyte-Klf1-ChIP-Seq(GSE20478)/Homer | 1e-2 | -6.442e+00 | 0.8010 | 5.0 | 11.90% | 989.7 | 2.04% | motif file (matrix) | svg |
| 3 | | Klf4(Zf)/mES-Klf4-ChIP-Seq(GSE11431)/Homer | 1e-2 | -6.357e+00 | 0.8010 | 8.0 | 19.05% | 2647.2 | 5.45% | motif file (matrix) | svg |
| 4 | | ATHB53(HB)/col-ATHB53-DAP-Seq(GSE60143)/Homer | 1e-2 | -5.085e+00 | 1.0000 | 4.0 | 9.52% | 850.4 | 1.75% | motif file (matrix) | svg |
| 5 | | EHF(ETS)/LoVo-EHF-ChIP-Seq(GSE49402)/Homer | 1e-2 | -5.022e+00 | 1.0000 | 11.0 | 26.19% | 5578.7 | 11.49% | motif file (matrix) | svg |
| 6 | | SPL15(SBP)/colamp-SPL15-DAP-Seq(GSE60143)/Homer | 1e-2 | -4.633e+00 | 1.0000 | 8.0 | 19.05% | 3521.2 | 7.25% | motif file (matrix) | svg |

**B**

| Rank | Motif | P-value | log P-pvalue | % of Targets | % of Background | STD(Bg STD) | Best Match/Details | Motif File |
|---|---|---|---|---|---|---|---|---|
| 1 | | 1e-14 | -3.376e+01 | 4.33% | 0.37% | 47.2bp (55.8bp) | Hand2(bHLH)/Mesoderm-Hand2-ChIP-Seq(GSE61475)/Homer(0.683) More Information \| Similar Motifs Found | motif file (matrix) |
| 2 | | 1e-14 | -3.375e+01 | 1.73% | 0.01% | 70.7bp (61.2bp) | RBPJ/MA1116.1/Jaspar(0.703) More Information \| Similar Motifs Found | motif file (matrix) |
| 3 | | 1e-14 | -3.354e+01 | 5.19% | 0.60% | 55.0bp (53.4bp) | ZmHOX2a(1)(HD-HOX)/Zea mays/AthaMap(0.649) More Information \| Similar Motifs Found | motif file (matrix) |
| 4 | | 1e-13 | -3.160e+01 | 1.52% | 0.01% | 17.6bp (52.9bp) | RPN4(MacIsaac)/Yeast(0.652) More Information \| Similar Motifs Found | motif file (matrix) |
| 5 | | 1e-12 | -2.956e+01 | 2.16% | 0.05% | 52.5bp (58.1bp) | WIP5(C2H2)/colamp-WIP5-DAP-Seq(GSE60143)/Homer(0.655) More Information \| Similar Motifs Found | motif file (matrix) |
| 6 | | 1e-12 | -2.821e+01 | 4.55% | 0.56% | 46.4bp (52.4bp) | br(var.3)/MA0012.1/Jaspar(0.765) More Information \| Similar Motifs Found | motif file (matrix) |
| 7 | | 1e-12 | -2.818e+01 | 1.08% | 0.00% | 23.8bp (19.4bp) | SNRNP70K(RRM)/Drosophila_melanogaster-RNCMPT00143-PBM/HughesRNA(0.764) More Information \| Similar Motifs Found | motif file (matrix) |
| 8 | | 1e-12 | -2.818e+01 | 1.08% | 0.00% | 44.9bp (0.0bp) | PH0166.1_Six6_2/Jaspar(0.679) More Information \| Similar Motifs Found | motif file (matrix) |
| 9 | | 1e-12 | -2.816e+01 | 2.16% | 0.06% | 40.7bp (49.0bp) | Pit1+1bp(Homeobox)/GCrat-Pit1-ChIP-Seq(GSE58009)/Homer(0.707) More Information \| Similar Motifs Found | motif file (matrix) |

**Supplemental Figure 12: a**, Known motif enrichment using HOMER. **b,** *de novo* motif enrichments using HOMER.

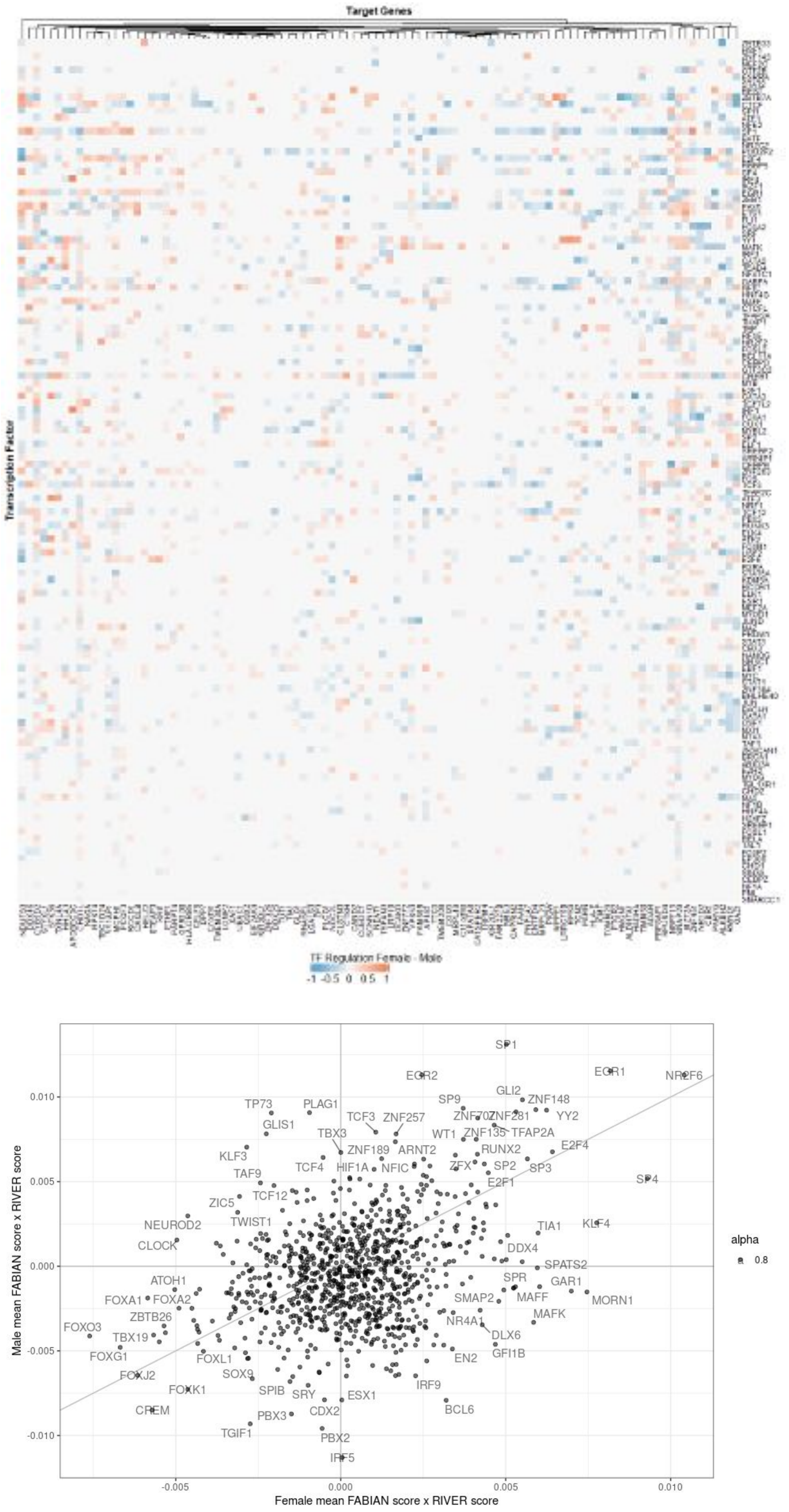


**Supplemental Figure 13: a**, For each TF-rare variant pair, FABIAN predicts a binding score. The maximum score for a given TF-rare variant pair was collapsed to the gene level. The rows are TFs and columns are genes. The values are the binding score x RIVER score, with the female score subtracted from the male score. **b,** The FABIAN x RIVER collapsed score for same genes in males vs females.

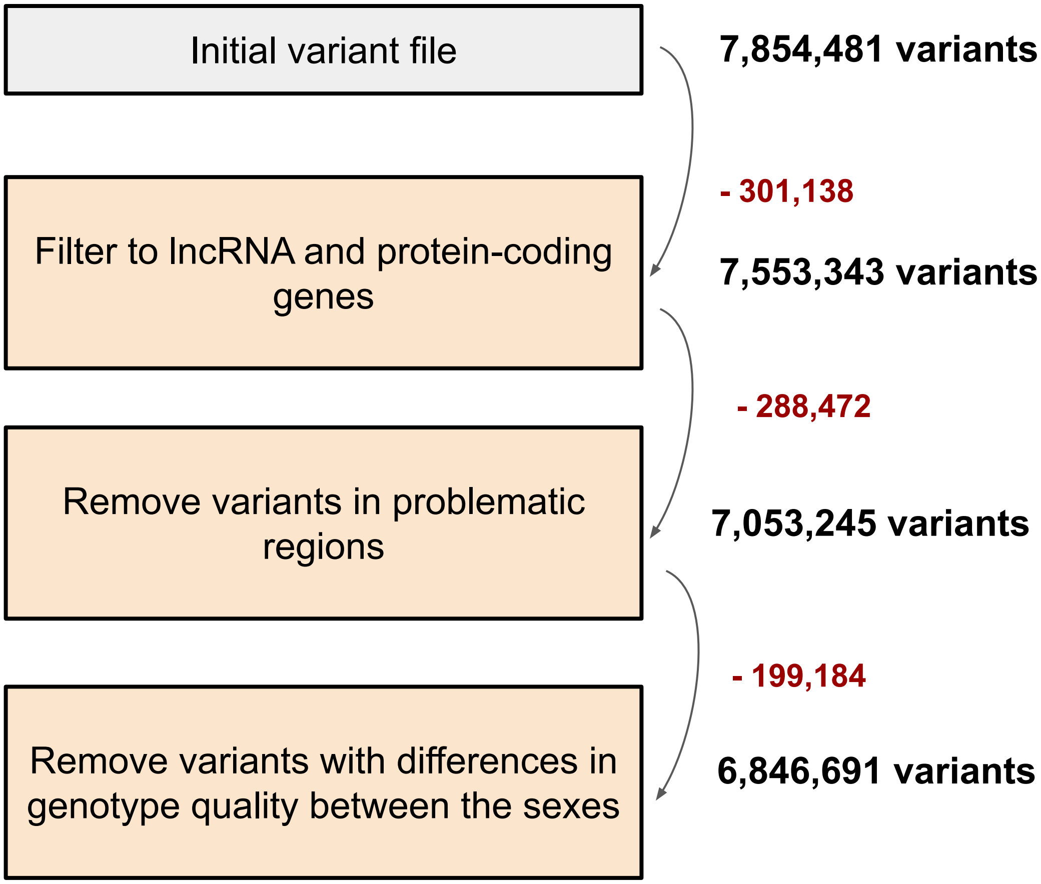


**Supplemental Figure 14: Variant filtration.** Variant filtration process. In red is the number of variants removed for each after each filtration process.

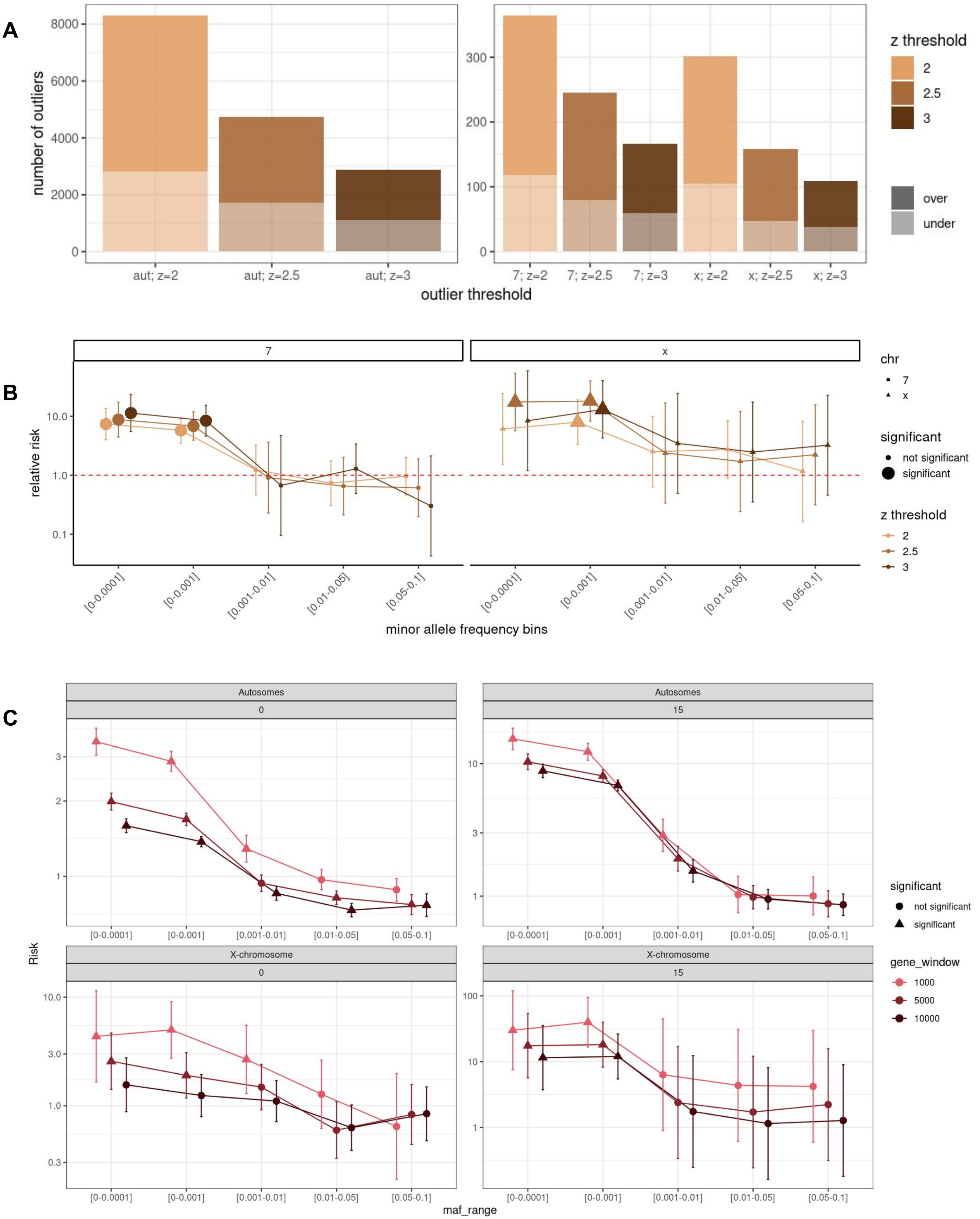
A
number of outliers
8000
6000
4000
2000
0
300
200
100
0
aut; z=2
aut; z=2.5
aut; z=3
7; z=2
7; z=2.5
7; z=3
x; z=2
x; z=2.5
x; z=3
outlier threshold
z threshold
2
2.5
3
over
under
B
7
x
relative risk
10.0
1.0
0.1
[0-0.0001]
[0-0.001]
[0.001-0.01]
[0.01-0.05]
[0.05-0.1]
minor allele frequency bins
chr
7
x
significant
not significant
significant
z threshold
2
2.5
3
C
Autosomes
0
Autosomes
15
X-chromosome
0
X-chromosome
15
Risk
3
2
1
10
3
1
10.0
3.0
1.0
0.3
100
10
1
[0-0.0001]
[0-0.001]
[0.001-0.01]
[0.01-0.05]
[0.05-0.1]
maf_range
significant
not significant
significant
gene_window
1000
5000
10000

**Supplemental Figure 15: Impact of parameterizations. a,** Impact on relative risk enrichments of the gene window which specifies how far away variants are to a given gene. This is done across the X-chromosome and autosomes, as well as a CADD threshold of zero and fifteen. **b,** The number of outliers that occur at different z-score thresholds across the autosomes, x-chromosome, and chromosome 7. **c**, The impact of z-score thresholds on relative risk enrichments.
